# Charting the nanotopography of inner hair cell synapses using MINFLUX nanoscopy

**DOI:** 10.1101/2025.04.22.649963

**Authors:** Rohan Kapoor, Hyojin Kim, Evelyn Garlick, Maria Augusta do R. B. F. Lima, Torben Ruhwedel, Wiebke Möbius, Fred Wolf, Tobias Moser

**Affiliations:** Institute for Auditory Neuroscience and InnerEarLab, University Medical Centre Göttingen, Göttingen, Germany; Auditory Neuroscience and Synaptic Nanophysiology Group, Max-Planck-Institute for Multidisciplinary Sciences, Göttingen, Germany; International Max Planck Research School (IMPRS) Molecular Biology, Göttingen Graduate School for Neuroscience and Molecular Biosciences (GGNB), Georg-August-Universität, Göttingen, Germany; Abberior Instruments, Hans-Adolf-Krebs-Weg 6, 37077 Göttingen, Germany; Electron Microscopy Unit, Department of Neurogenetics, Max Planck Institute for Multidisciplinary Sciences, Göttingen, Germany; Multiscale Bioimaging Cluster of Excellence, MBExC, Göttingen, Georg-August-Universität, Germany; Campus Institute for Dynamics of Biological Networks, Georg-August-Universität, Göttingen, Germany

## Abstract

For us to hear, the cochlea encodes sounds into neural signals at synapses of inner hair cells (IHCs) and the auditory nerve with remarkable fidelity. To achieve the high rates of temporally precise synaptic transmission over long periods of time, IHCs employ sophisticated ribbon-type active zones (AZ). In order for us to understand synaptic sound encoding, we need to decipher the underpinning molecular topography of these synapse which had remained challenging due to technological limitations. Here we applied 3-dimensional minimal flux optical nanoscopy to mouse IHC synapses to chart the position of key pre- and postsynaptic proteins with single digit nanometre resolution of imaging. We demonstrate that nanoclusters of channels and interacting proteins govern the topography of AZs and postsynaptic densities (PSDs). We count synaptic proteins, their nanoclusters and determine their spatial organization feeding into computational modelling of AZ function. In conclusion, this study reveals a nanocluster-based molecular AZ and PSD topography, likely serving as functional modules in synaptic sound encoding.

## Introduction

Inner hair cells (IHCs) of the cochlea, like retinal photoreceptors, form specialized synapses that are controlled by graded receptor potentials, providing greater bandwidth for relaying sensory information than achievable by all-or-none action potentials (for review see *1*). The hallmark of these synapses are the eponymous synaptic ribbons – electron dense structures that tether synaptic vesicles (SVs) in vicinity of the presynaptic active zone (AZ) which enables fast, precise and tireless neurotransmission at these synapses. Synaptic ribbons represent the most elaborate specializations among other presynaptic electron dense projections that tether SVs, which include the small *presynaptic grids* found at vertebrate central nervous system (CNS) synapses and *T bars* in *Drosophila* (for review see (*2*). IHC synaptic ribbons are typically ellipsoidal, round or wedge-like in shape, ranging between 100 and 500 nm along their maximal extent, about three times smaller than in retinal rod photoreceptors (*3*). The protein RIBEYE is the core component of all ribbons (*4–7*) and has been proposed to self-assemble into a multi-lamellar structure which constitutes the ribbon scaffold (*8*), along with the protein Piccolino (*9–11*).

Over the decades, electron microscopy and tomography have been extensively used to study the architecture of these specialized synapses, in particular shedding light on the abundance, arrangement and tethering of SVs (*12–17*). On the other hand, light microscopy, in particular super- resolution STED microscopy, has provided high resolution insights into synaptic assemblies of key molecular components, including AZ proteins like Bassoon (*14*, *18*), RIM2 (*19*), RIM-BP2 (*20*) and voltage-gated Ca_V_1.3 Ca^2+^ channels (*18*, *21*). Ca_V_1.3 channels have been shown to cluster underneath the presynaptic AZ, forming an on average a 400 nm long narrow stripe at 60% of the synapses (*6*, *11*, *18*, *20*).

However, methods used thus far could not answer key remaining questions: How does the arrangement of Ca_V_1.3 channels relate to vesicular release sites? How is the topography of Ca_V_1.3 channels and AZ components: e.g. are the half-micrometre-long clusters of Ca_V_1.3 channels and other AZ proteins (“microclusters”) further composed of smaller “nanoclusters”? Such nanoclusters were shown by STORM imaging for the multidomain AZ protein Munc13-1 at conventional CNS synapses and were interpreted as molecular correlates of SV release sites (*22*). Moreover, studies at CNS synapses have shown nanoclusters of presynaptic AZ components to be tightly aligned with postsynaptic scaffold proteins and receptor nanoclusters suggesting trans-synaptic “nanocolumns” which may contribute to synaptic efficiency at such synapses (*23*). Finally, how RIBEYE and Piccolino assemble within the synaptic ribbon remains an important open question.

Several arrangements of Ca_V_ channels and vesicular release sites have been suggested for different synapses, including random positioning of Ca^2+^ channels and SVs (*24*), Ca^2+^ channels forming a ring around SVs (*25*), positioning of SVs at the perimeter of Ca^2+^ channel clusters (*26*, *27*), a one- to-one stoichiometry of SV and Ca_V_ channel (*28–33*), or an exclusion zone model where Ca^2+^ channels do not cluster but are simply excluded from a 50 nm zone around docked SVs (*34*). A paradigmatic topography of Ca^2+^ channels and SV release sites supporting one-to-one stoichiometry is the longitudinal array of functional units suggested based on studies of the neuromuscular junction (*30*) and the rod ribbon synapse (*33*). SV release at mature IHC ribbon synapses seems to be governed by 1-3 Ca^2+^ channels exerting a Ca^2+^ nanodomain-like control (*14*, *15*, *21*, *35–39*). The effective coupling distance between Ca^2+^ channels and the Ca^2+^ sensor for SV exocytosis at the IHC AZ has been estimated to be within 17 nm (*37*). This results in a near linear dependence of release on the number of open Ca_V_ channels offering a wide dynamic range of the synaptic transfer function that tracks the open probability of Ca_V_1.3 Ca^2+^ channels for optimal control of sound intensity coding by the IHC receptor potential (e.g. (*38–40*). Moreover, a tight channel-vesicle coupling places the low affinity Ca^2+^ sensor of exocytosis (*41*), most likely Otoferlin (*42–44*) into the saturating range thereby reducing synaptic delay and its variance for fast and temporally precise neurotransmission (*40*). Biophysical modelling indicates that a subset of Ca^2+^ channels at the periphery of the typically stripe-like microcluster of the AZ exert a Ca^2+^ nanodomain control akin to “private channels” of the release site (*21*, *37*). However, all this remains inference from indirect approaches and the precise topography of Ca^2+^ channels and SV release sites remains to be elucidated.

Advances in MINFLUX nanoscopy (*45*, *46*) have now made it possible to investigate the spatial arrangement of multiple proteins with fluorescence microscopy at a molecular-level spatial resolution in 3D, offering to bridge conventional electron and light microscopy techniques with molecular specificity. Despite this, routine application of MINFLUX has typically used cultured cells and its utility has remained more limited for native tissues. This is primarily because of (i) high sensitivity of MINFLUX to photons emerging from background signal, which is far more pronounced in tissue samples owing to autofluorescence caused by chemical fixation, out-of-focus signal, and general “stickiness” of tissue samples to adsorb primary and secondary antibodies non-specifically; and (ii) spherical aberrations and refractive index mismatch in the tissue, which progressively worsens imaging conditions if the structure of interest is away from the coverslip and deeper into the tissue. Here, we have established MINFLUX imaging of 1 μm semithin sections from Epon embedded mouse cochleae which were etched prior to immunostaining. This allowed for convenient, reproducible and high-yielding tissue sample preparation for MINFLUX, with synaptic structures immobilized close to the coverslip allowing minimal spherical aberrations and fluorescence background.

Using MINFLUX imaging of synapses in such semithin sections, we have mapped the distribution of proteins of the synaptic ribbon (RIBEYE, Piccolino), of the AZ proper (Ca_V_1.3 and RBP2, Rab3 interacting molecule Binding Protein 2) and the postsynapse (Homer1 and GluA2) to reveal their 3D spatial arrangement. We further quantified the data to deduce “molecular counts” of synaptic proteins and we observed that the spatial arrangement of most of these proteins follows a nanocluster-based molecular organisation. Biophysical modelling indicates that such a nanocluster-based organisation may support the efficiency of synaptic transmission at IHC ribbon synapses.

## Results

### Immunolabeled 1μm sections for MINFLUX imaging of cochlear synapses

MINFLUX allows rapid imaging at molecular level spatial resolution (1 - 3 nm). Tissues of varying thicknesses and complexities do not match well with the high sensitivity of single molecule localizations due to background fluorescence and spherical aberrations. Recently MINFLUX was used to study the nanoarchitecture of the photoreceptor ribbon synapse by developing heat assisted rapid dehydration (HARD), which allows immobilization of a thin layer of retinal cells on the coverslip (*33*). However, the tissue complexity of the cochlea with its osseous labyrinth, spacious cavities and delicate organ of Corti, the sensory epithelium, made it challenging for us to adapt HARD for studying hair cell synapses that often were at a distance > 3 μm from the coverslip (Fig. S1A-D). We also attempted using cochlear cryosections of 10 – 16 μm thickness (Fig. S1E-I). Treating the samples using 0.1% NaBH4 solution or 0.1M tris(hydroxymethyl)aminomethane (Tris) prior to blocking and immunostaining, drastically quenched tissue autofluorescence and allowed us to obtain an improved signal to noise ratio of immunolabelled ribbon synapses. While we could perform preliminary MINFLUX imaging in such sections, the quality still remained inadequate, owing to high photon emissions from the tissue background. We refrained from using paraffin-embedded sections which often have limited epitope availability for antibody binding.

Instead, several previous studies have shown how embedding tissue samples in epoxy resin followed by sectioning and etching the resin can help overcome this problem and allow nanometre sections with largely preserved labelling efficiency (*47–49*). We used sections of 0.3 – 1 μm thickness from chemically fixed, decalcified cochleae which we embedded in Epon (see Fig. 1A). After sectioning, we removed Epon prior to immunostaining by etching the samples with NaOH in absolute ethanol (*50*). Such samples were highly appropriate for MINFLUX imaging as the minimal thickness of the tissue ensures low background, high signal quality, and the synapses were typically close to the coverslip. One μm thick sections turned out to be most suitable with homogenous antibody penetration and full enclosure of synapses, whereas thinner sections of 300 and 600 nm often only featured partial synaptic structures. Advantages of this sample preparation approach are simplicity, high reproducibility and high yield of hundreds of sections from a single cochlea. We could obtain good quality staining for a majority of IHC markers, AZ proteins, synaptic ribbon markers and postsynaptic proteins as summarized in Fig. S2A, with the exception of AZ protein Bassoon and the putative Ca^2+^ sensor of synaptic vesicle fusion Otoferlin, for which we could obtain only faint signals with substantial background. Staining IHCs with antibodies directed against Calretinin or VGLUT3 allowed easy identification of IHCs in tissue sections for imaging (Fig. 1B). We also compared the immunofluorescence intensity of synaptic ribbons in these samples with those in acute preparations of the organ of Corti stained with the same antibody concentrations and imaged in parallel with the same laser power. We did not find any reduction in immunofluorescence intensity, implying that the Epon embedding and etching likely do not contribute to loss of epitopes, (Fig. S2B). Not unexpected (*20*, *35*), stainings for Ca_V_1.3 Ca^2+^ channels and RBP2 were not optimal with 4% formalin fixation, but improved considerably with glyoxal fixation (*51*) and could then be used for MINFLUX imaging. In summary, we propose that using semithin sections of Epon embedded tissue can provide a simple and highly adaptable method for routinely performing MINFLUX imaging, prospectively also on different tissue types.

**Fig. 1.**
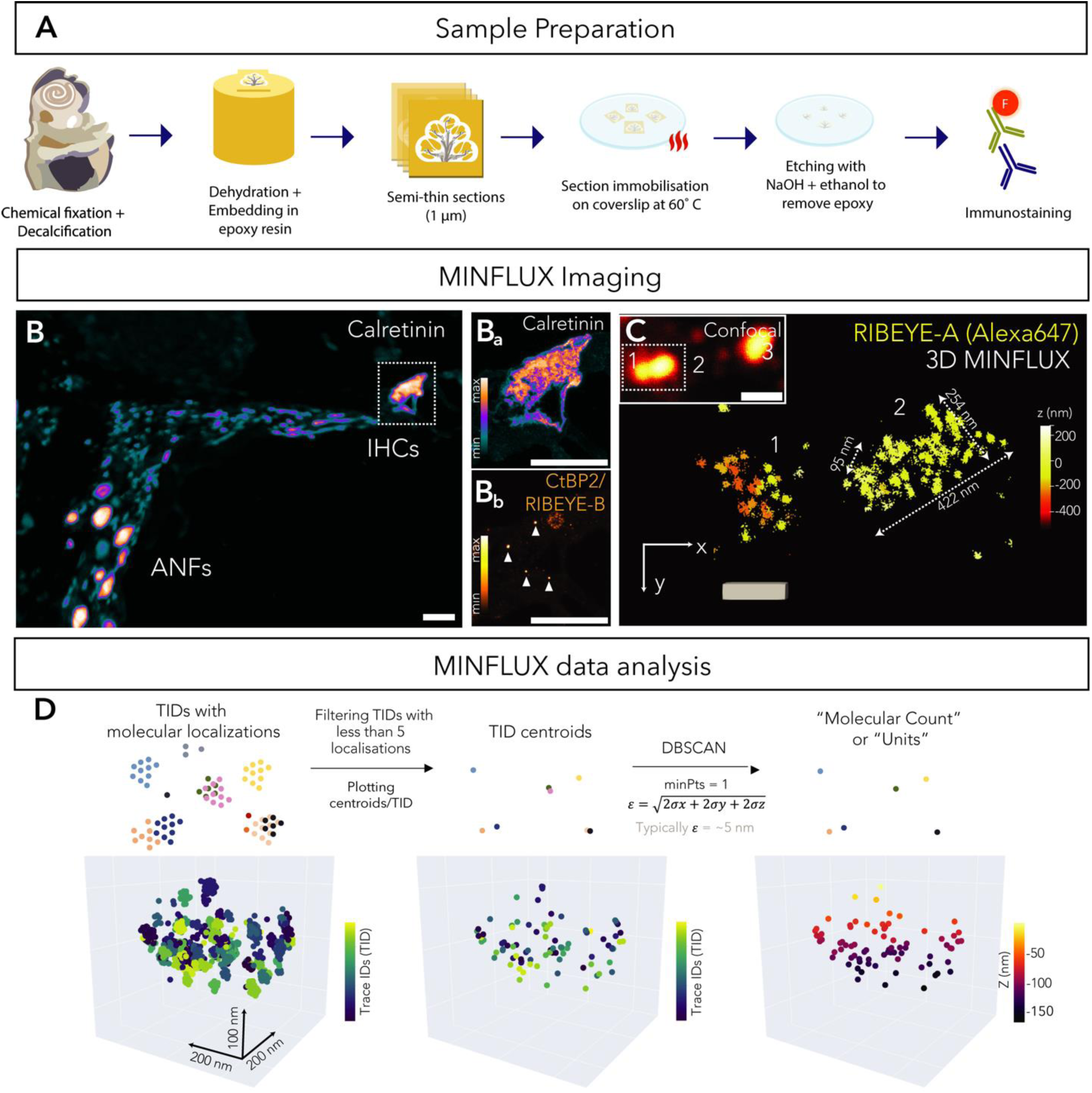
MINFLUX sample preparation, imaging and data analysis for cochlear tissue. (A) Schematic illustrating preparation of 1 μm semithin cochlear sections used for MINFLUX imaging **(B)** Representative confocal image (maximum intensity projection) of 1 μm cochlear section labelled with an antibody against Calretinin (labelling IHCs and auditory nerve fibres (ANFs)). Panels **(Ba)** and **(Bb)** show zoom-in of IHC with individual channels representing Calretinin and CtBP2/RIBEYE (orange, labelling synaptic ribbons, marked with arrows) stainings. Scale bar = 20 μm; individual channels shown with intensity-coded look up tables. **(C)** Exemplary images of synaptic ribbons acquired using 3D MINFLUX. An antibody against RIBEYE A-domain was used. Inset shows corresponding confocal image (single plane) of the ribbons. Note how the two ribbons (labelled with numbers) appear as one entity in the confocal image due to limited resolution. Colour code represents the distances along the z-axis. Scale bar = 200 nm. **(D)** Illustration (upper panel, not to scale) and exemplary 3D MINFLUX data of a synaptic ribbon (lower panel) depicting the analysis scheme. Different colours in the upper panel represent individual Trace IDs (TIDs), points represent molecular localizations belonging to a TID. We first determined the centroid of each TID, rejected TIDs with less than 5 localizations, and then performed clustering using DBSCAN. The clusters so obtained are referred to as “molecular counts” or “units” and correspond to centroids from different TIDs at a distance more than ∼2σ in each dimension.

### 3D MINFLUX reveals IHC ribbons packed with RIBEYE molecules

We started with 3D MINFLUX imaging of the synaptic ribbon, done here for the first time to our knowledge. We used a primary antibody labelling RIBEYE, the core scaffold protein of the ribbon, in 1 μm cochlear sections from 2-week-old C57BL6/J mice, followed by a secondary antibody conjugated to Alexa647. We found the raw localization precision of the imaging to be between 3.9 and 5.9 nm in all three dimensions. Synaptic ribbons were ellipsoidal in shape, and on average about 400 nm in length, 150 nm in width and 200 nm in height (measures of an exemplary ribbon depicted in Fig 1C). The topography of RIBEYE from MINFLUX data did not seem to indicate any regular, lamellar pattern of arrangement as has been reported to occasionally occur in electron micrographs of mature synaptic ribbons (*3*, *52*, *53*). We also encountered double ribbons seemingly from the same synapse as shown in Fig. 1C, which appear as one entity using confocal imaging but could be resolved well into two discernible synaptic ribbons using MINFLUX. Three of the six imaged ribbons appeared to have a somewhat hollow centre potentially corresponding to the translucent cores in otherwise electron-dense IHC ribbons of adult mice found in previous EM studies (*11*, *52*, *54–56*). Alternatively, the finding might reflect reduced penetration of the antibody into a densely packed ribbon.

For quantification of the MINFLUX data, we used a custom-written analysis pipeline employing the Density-Based Spatial Clustering of Applications (DBSCAN) clustering algorithm. The localization of an emitting molecule is typically performed by the microscope several times by repetition of the last two MINFLUX iterations. Since these successive localizations belong to the same blinking event or localization “burst”, they are assigned the same trace ID (TID). We used MINFLUX images filtered for quality parameters (see Methods section) and determined the centroid of each TID (for TIDs with at least 5 localizations) by averaging coordinates of all localizations per TID. We then implemented DBSCAN clustering with minimum number of points = 1 and radius e of cluster defined by the square root of sum of 2σ in each dimension (∼5 nm) to obtain “molecular counts” or “units” (*57*); see Fig. 1D for schematic representation. These molecular counts represent the number of emitting molecules emerging from different blinking events that may localize very close to each other, while minimizing the contribution from repeated localizations. Synaptic ribbons appeared to be packed with on average 103 ± 19 RIBEYE molecules/synapse (S.D. = 47, range = 45 – 159, N = 6 synapses); average density per unit volume = 21458.3 molecules/µm^3^, which is substantially lower than prior estimates for retinal bipolar neurons of goldfish obtained from peptide labelling (∼4000 RIBEYE molecules/ribbon, (*58*)). This might be largely due to technical differences, as the ribbon dimensions are similar (*1*, *59*).

### Piccolino forms an arch around the membrane-distal synaptic ribbon

Piccolino (*9*) is a short ribbon-synapse specific splice-variant of the multi-domain AZ protein Piccolo, found in the cochlea and the retina. Piccolino appears to be distributed exclusively at the ribbon (*10*, *11*, *54*, *60*, *61*) and has been shown to interact with RIBEYE directly (*10*). We immunostained 1 μm semi-thin cochlear sections from p15 C57BL6/J mice with VGLUT3 (Alexa488, to label IHCs), CtBP2/RIBEYE (Alexa541, labelling synaptic ribbons) and Piccolino (Alexa647), using an antibody that labels Piccolino but also full-length Piccolo. Our attempts to perform two-colour MINFLUX for Piccolino and CtBP2/RIBEYE did not work out well and spectral separation of signals from the two proteins was unclear. To distinguish Piccolino at afferent ribbon synapses from full-length Piccolo at efferent synapses, formed by lateral olivocochlear neurons onto SGN boutons, we imaged only Piccolino immunofluorescent puncta that colocalised with CtBP2/RIBEYE immunofluorescence observed at confocal resolution. 3D MINFLUX revealed Piccolino to be distributed over the synaptic ribbon in the form of an arch, presumably enveloping the membrane-distal (apical) circumference of the ribbon (Fig. S3A, B) with approximate dimensions 400, 120 and 200 nm along length, width and height respectively. This is also in agreement with previous immunogold EM and STED microscopy data which have shown Piccolino to be similarly distributed along the IHC ribbon, being particularly enriched at the apical end of the ribbons of ∼p14 mice and being largely absent at the very base of the ribbon near the AZ (*11*, *54*).

### RBP2 and Ca_V_1.3 are arranged in a narrow stripe under the ribbon

As for Piccolino, we imaged only RBP2 immunoreactive puncta that colocalised with synaptic ribbons (Fig. 2A, B) as RBP2 also localises at adjacent efferent synapses (*20*). 3D MINFLUX images show RBP2 to be arranged in a linear stripe-like manner (on average around 382 nm in length, 94 nm in width and 60 nm in height, N = 10 synapses) underneath the synaptic ribbon, along its length (Fig. 2C, D, E) in agreement with published STED microscopy data (*20*). We also performed 3D MINFLUX imaging for the AZ protein RIM2, which also yielded a similar linear, stripe-like arrangement of the protein underneath the ribbon, with some synapses showing a more complex organization (Fig. S4). Unfortunately, our RIM2 stainings gave results only occasionally and hence this data was not used for quantitative analysis (N = 4 successfully imaged synapses).

**Fig. 2.**
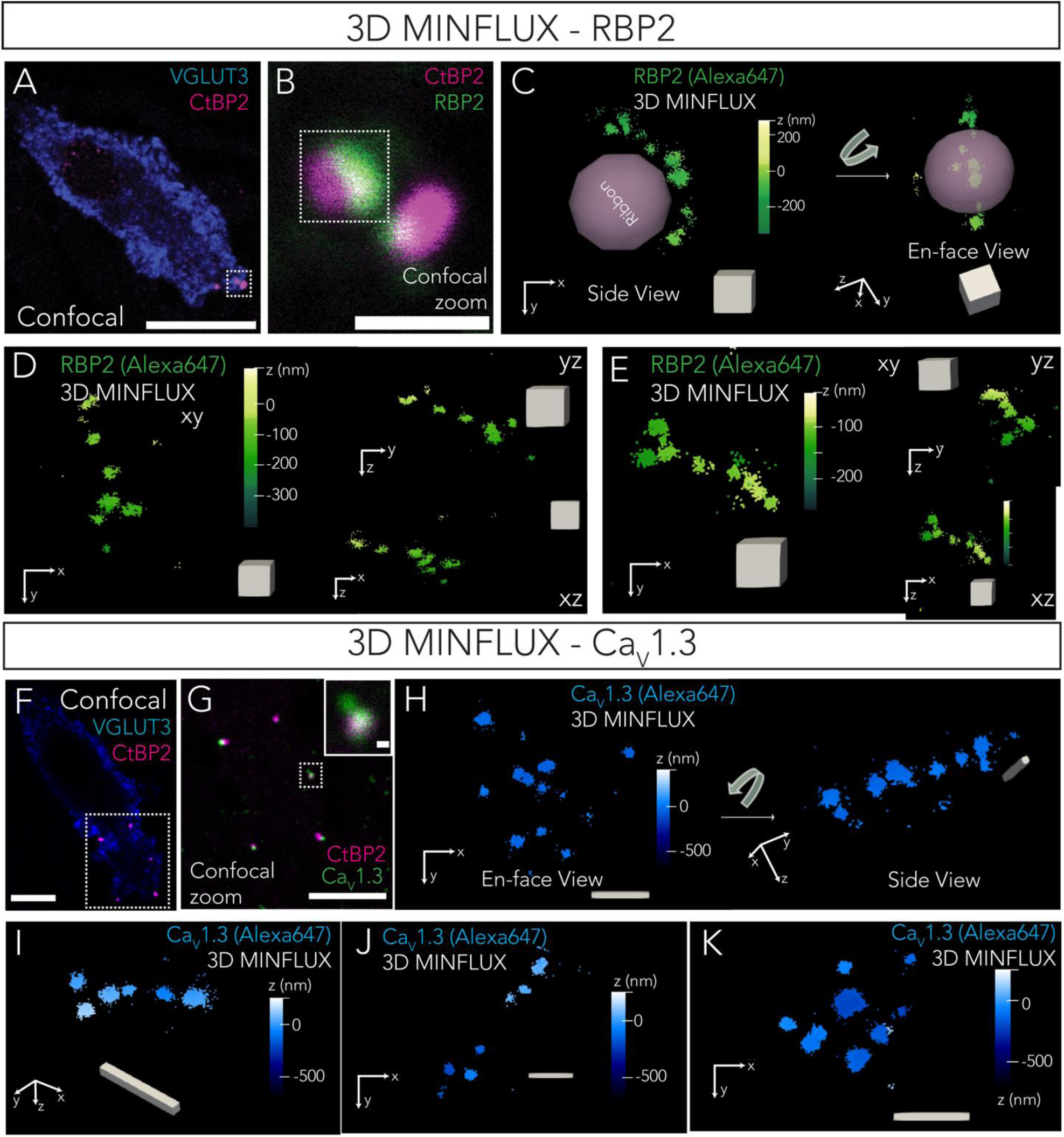
3D MINFLUX imaging of the AZ proper – RBP2 and CaV1.3. **(A)** Confocal overview of IHC used for imaging (VGLUT3, blue) and the location of the ribbon synapses (CtBP2/RIBEYE, magenta). Scale bar = 10 µm. **(B)** Zoom-in of the ribbon synapses. Here the corresponding RBP2 puncta have been imaged (green). Scale bar = 1 µm. **(C)** Raw 3D MINFLUX imaging of RBP2 from the synapse marked in (B) shows the spatial distribution of RBP2 along a linear AZ. The pink ovoid shown depicts the presumed position of the ribbon based on confocal two-colour imaging. **(D, E)** Further exemplary 3D MINFLUX of RBP2 highlight the line-shaped morphology of the AZ; cube shown for scale in (C, D, E) has edge of 100 nm **(F)** Confocal overview of the IHC used for imaging (VGLUT3, blue) and the location of ribbon synapses (CtBP2/RIBEYE, magenta). Scale bar = 5 µm. **(G)** Confocal zoom-in of the ribbon synapses showing colocalising CaV1.3 labelling (green). Scale bar = 5 µm. **(H)** Raw 3D MINFLUX images of CaV1.3 channels from the synapse shown in inset of (G). From the top, CaV1.3 shows a complex spatial arrangement. Rotating the image as shown gives a linear arrangement of the localisations along a side view of the AZ. **(I, J, K)** Further exemplary 3D MINFLUX images of CaV1.3 channels. Note the double-line shaped arrangement of CaV1.3 channels in (K). Scale bar for (I, J, K) is 200 nm. In all MINFLUX images, colour code represents distances along z-axis.

Like RBP2, Ca_V_1.3 localizations typically appeared to be distributed in narrow stripes (Fig. 2F-K), in agreement with previous observations from STED imaging where over 60% of Ca_V_1.3 clusters take this shape (*6*, *11*, *14*, *18*, *20*). Ca_V_1.3 channels viewed *en face* at eight of the eighteen synapses imaged resembled a more complex and wider arrangement of channels (as shown in STED images before (*18*)). However, on rotating, these typically also appeared linear (see Fig. 2H, another exemplary synapse shown in Fig. S5A), likely representing a side view along the plasma membrane. In one particular synapse of these, we also encountered Ca^2+^ channels seemingly arranged in two lines (akin to double-line shaped Ca_V_1.3 clusters seen in STED images (*18*)) as shown in Fig. 2K. The distance between the two lines was around 160 nm from centre points. On the other hand, another eight of the eighteen synapses imaged showed a more defined linear morphology (Fig. 2I, J) in all perspectives, irrespective of the orientation in which they were observed (see Fig. S5B). These linear stripes of channels on average appeared 422 nm long, 102 nm wide and 47 nm high, and presumably run underneath the synaptic ribbon along its length (∼400 nm). The line appeared thicker along one side than the other in half of these synapses. The remaining two of the eighteen synapses seemed reminiscent of “spot-like” Ca_V_1.3 channel clusters reported previously (*18*) and appeared to have highly concentrated localisations in two-three “spots” with no seemingly defined morphology (Fig. S5C). A summary of all imaged CaV1.3 clusters, their heterogenous morphologies and calculated parameters has been provided in Table S5.

We estimated between 9 – 70 Ca_V_1.3 molecular counts per synapse (on average 35 ± 4 counts/synapse, S.D. = 18, median = 34, N = 18 synapses). These estimates fall within limits of previous functional estimates (20 – 330 Ca^2+^ channels/AZ) but are lower than the median counts (60 – 120 Ca^2+^ channels/AZ (*18*) indicating an underestimation by MINFLUX e.g. due to incomplete labelling and/or detection. Indeed, we assume a labelling efficiency of ∼55 – 60% similar to what was reported for immunoelectron microscopy studies of AMPA receptors (*62*). RBP2 interacts with voltage-gated Ca^2+^ channels (*63*) and at the IHC ribbon synapse, it has been shown to play a role in promoting larger complements of Ca_V_1.3 channels (*20*). Not surprisingly, RBP2 counts appear rather similar to counts of Ca_V_1.3 channels on average (30 ± 3 counts/synapse, S.D. = 9, range = 20 – 43, N = 10 synapses). As above, we reason that it is likely that our estimates provide a lower bound of the number of RBP2 molecules on account of not all epitopes being available for antibody binding.

### Postsynaptic AMPA receptors are organised in a ring engulfing the presynapse

3D MINFLUX imaging of Homer1 shows that the postsynaptic density of the afferent synapse is packed with hundreds of Homer1 molecules (Fig. 3A-D). The top view reveals dense localizations spread across the postsynaptic density, with intermediate regions of relatively sparse localizations that give the impression that the Homer1 scaffold may be arranged in flat, roundish nanoclusters. Side views of the 3D MINFLUX images shows the characteristic flat, plate-like postsynaptic density (Fig. 3D). The postsynaptic density appeared to be elongated along one axis (∼800 nm) and shorter (∼300 nm) along the other axis and appeared relatively flat along its height (∼100 nm). We also performed two-colour 2D MINFLUX by labelling synaptic ribbons and postsynaptic Homer1. For two-colour MINFLUX imaging, the fluorophores were excited with the same excitation wavelength and two different spectral detectors were used to detect emissions which could be separated via spectral unmixing, as described in the methods section. Fig. 3E shows a side-profile of a synaptic ribbon seated atop of a postsynaptic density with the synaptic cleft in between (distance = ∼65 nm from the base of ribbon to the top of the postsynaptic density).

**Fig. 3.**
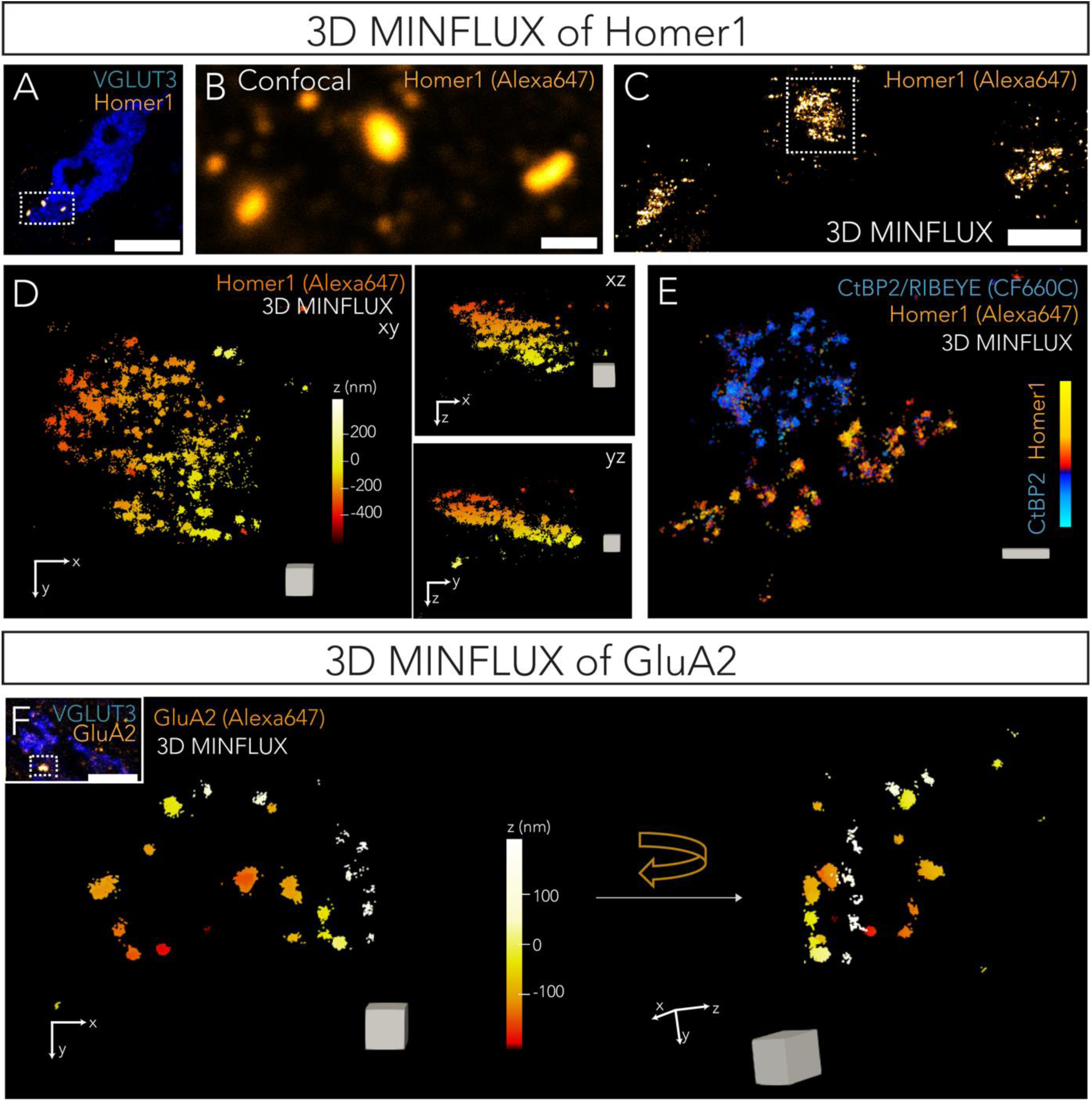
3D MINFLUX imaging of postsynaptic Homer1 and AMPA receptors. **(A)** Representative confocal overview of an IHC (single plane) labelled with an antibody against VGLUT3 (blue). Postsynaptic SGN boutons are labelled with an antibody against Homer1 (orange); scale bar = 10 µm. **(B)** Confocal zoom-in from (A) showing three postsynaptic densities, with the VGLUT3 channel removed for clarity; scale bar = 1 µm. **(C)** 3D MINFLUX z-projection of Homer1 showing an overview of the three postsynaptic densities in (A). Colour code represents number of localisations/voxel; scale bar = 1 µm. **(D)** Raw 3D MINFLUX image displaying a closer view of the postsynaptic density highlighted in (C). Colour code represents distances along the z-axis. Panels on the right show y- and x- projections (from top to bottom) respectively. Cube shown for scale has an edge of 100 nm. **(E)** Two-colour 2-D MINFLUX images of a ribbon synapse with the presynapse marked by labelling against CtBP2 and postsynapse marked by Homer1 labelling. Scale bar = 100 nm. **(F)** Raw 3D MINFLUX image of postsynaptic AMPA receptors (labelled with an antibody against GluA2) shows a ring-like arrangement of the AMPA receptors localisations at the postsynapse. Inset shows a confocal overview of the synapse imaged, scale bar = 5 µm. Right panel shows the glutamate receptor ring turned clockwise for a side perspective (see orientation axis). Colour code represents distances along the z-axis, cube shown for scale has an edge of length = 100 nm.

Next, we investigated the topography of postsynaptic AMPA receptors in SGN boutons. Postsynaptic currents in SGNs are primarily believed to be driven by AMPA receptors and not NMDA or Kainate receptors (*64*) and SGN boutons have been shown to express mostly GluA2, along with GluA3 and GluA4 subunits of AMPA receptors, but not GluA1 (*65*). Using 3D MINFLUX, we imaged GluA2 localizations and found GluA2 in *en-face* synapses (3 of 8 synapses) to be arranged in a characteristic ring-like manner (Fig. 3F) as also shown previously with STED imaging (*66*, *67*) and immune-electron microscopy (*65*). In the remaining synapses, the ring-like arrangement of GluA2 was less clear as also shown before with STED imaging (*66*, *67*). The peak-to-peak distance was on average approximately 790 nm (from N = 8 synapses imaged *en face*), close to the long axis from Homer1 MINFLUX images.

We report on average 51 ± 12 GluA2 units/synapse (S.D. = 34, range = 24 – 110 from N = 8 synapses) which correspond well with estimates of AMPA receptors (∼60 particles/postsynaptic density) from immuno-gold EM data from guinea pig IHC ribbon synapses (*65*).

A complete summary of molecular counts, nearest neighbour distances between all molecular counts and volume of alpha shape fits has been provided in Table 1 and Fig. 4A-D for all the described pre- and post-synaptic proteins.

**Fig. 4.**
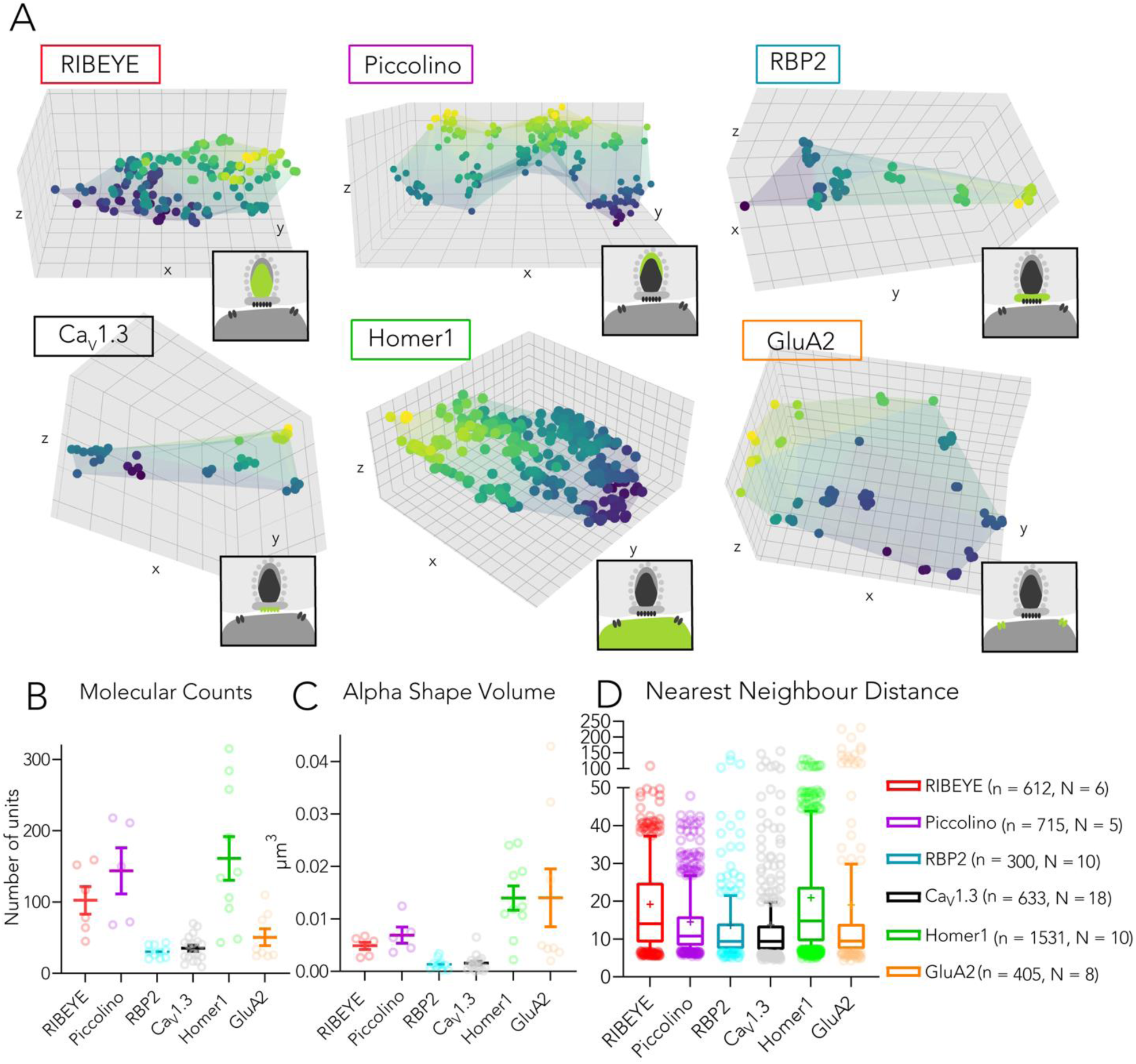
Mapping the topography of pre- and postsynaptic proteins at the IHC ribbon synapse. **(A)** Representative 3D plots showing individual molecular counts per synapse obtained from 3D MINFLUX images of RIBEYE, Piccolino, RBP2, CaV1.3, Homer1 and GluA2, along with superimposed convex hull rendering (translucent alpha shape fits). Colour scale is indicative of distances along the z-axis and scale of grid lines along all axes is 50 nm. Cartoon insets depict the localisation of the respective proteins at the ribbon synapse. **(B)** Plot showing mean ± SEM molecular counts from 3D MINFLUX images of different synaptic proteins. Data points represent individual synapses. **(C)** Plot depicting mean ± SEM of volume from alpha shape fits on coordinates of units. Data points represent individual synapses. **(D)** Box-whisker plot showing nearest neighbour distances for all units from different synaptic proteins. Individual data points represent molecular counts crosses represent mean values, central band indicates the median, whiskers represent 90/10 percentiles and boxes represent 75/25 percentiles.

**Table 1.** Summary of quantitative analysis of pre- and post-synaptic marker proteins. Molecular counts, nearest neighbour distances (NND) between all molecular counts (“units”) and the volume estimates from convex hull fits for different synaptic proteins have been provided.

|  | **RIBEYE** | **Piccolino** | **RBP2** | **Cav1.3** | **Homer1** | **GluA2** |
| --- | --- | --- | --- | --- | --- | --- |
| **Molecular counts/synapse (Mean, N)** | 103 | 144 | 30 | 35 | 161 | 51 |
| **SEM** | 19 | 32 | 3 | 4 | 31 | 12 |
| **SD** | 47 | 72 | 9 | 18 | 97 | 34 |
| **Median** | 98 | 150 | 30 | 34 | 138 | 31 |
| **Range** | 45 - 159 | 68 - 218 | 20 - 43 | 9 - 70 | 43 - 315 | 24 - 110 |
| **NND b/w all molecular counts (Mean, nm)** | 19.17 | 14.50 | 13.62 | 13.74 | 20.92 | 19.01 |
| **SEM** | 0.59 | 0.42 | 0.88 | 0.64 | 0.46 | 1.54 |
| **SD** | 14.59 | 11.11 | 15.34 | 16.03 | 18.15 | 30.99 |
| **Median** | 14.02 | 10.80 | 9.38 | 9.35 | 14.78 | 9.41 |
| **Volume of Alpha Shape (Mean, $\mu\text{m}^3$)** | 0.0048 | 0.0069 | 0.0013 | 0.0016 | 0.0140 | 0.0140 |
| **SEM** | 0.0007 | 0.0015 | 0.0003 | 0.0004 | 0.0023 | 0.0055 |
| **SD** | 0.0017 | 0.0035 | 0.0010 | 0.0015 | 0.0074 | 0.0156 |
| **Mean Density (counts/ $\mu\text{m}^3$)** | 21458.3 | 20869.6 | 23076.9 | 21875.0 | 11500.0 | 3642.8 |
| **N (No. of synapses)** | 6 | 5 | 10 | 18 | 10 | 8 |
| **n (Total molecular counts )** | 612 | 715 | 300 | 633 | 1531 | 405 |

### Two-colour-MINFLUX reveals distribution of VGLUT3-positive vesicular compartments around the synaptic ribbon

Next, we performed two-colour MINFLUX imaging by labelling synaptic ribbons with an antibody against CtBP2/RIBEYE (Alexa647 conjugated secondary) and SVs with VGLUT3 (CF660C conjugated secondary antibody), the vesicular glutamate transporter isoform found in inner hair cells (*68–70*). We observed a very dense labelling for VGLUT3 around the synaptic ribbon and also throughout the IHC (Fig. 5A, B; N = 4 MINFLUX images). The VGLUT3 signal appeared highly specific to IHCs with negligible signal outside the cell. IHCs have been shown to be densely packed with glutamatergic synaptic vesicles as demonstrated by the strong VGLUT3 immunofluorescence and colocalised Otoferlin immunofluorescence (*71*, *72*). VGLUT3 localizations may therefore reflect putative SVs around the synaptic ribbon and other VGLUT3- positive compartments in the cell. We performed cluster analysis of VGLUT3 units using DBSCAN with minimum number of VGLUT3 molecules = 6 (assuming 60% labelling efficiency, estimates suggesting 9-10 VGLUT1/2 per SV; *85*) and e = 20 nm (Fig. 5C). VGLUT3 units assigned into clusters were fitted with circles to estimate diameters of the putative vesicles which was 40.31 nm (median), and on average 44.30 ± 2.13 nm (S.D. = 17.42); Fig. 5D.

**Fig. 5.**
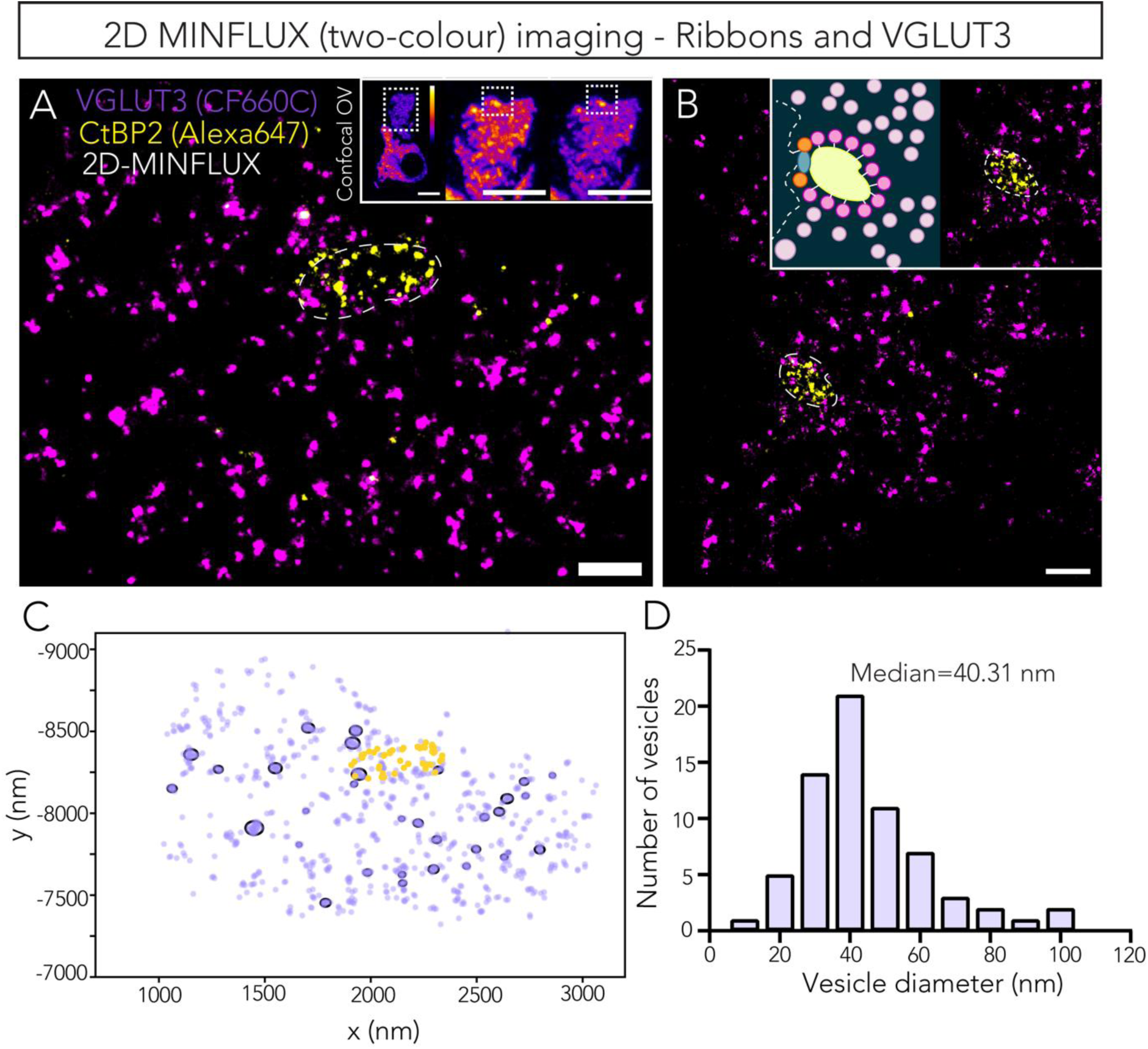
MINFLUX shows distribution of VGLUT3-positive membranes around the synaptic ribbon. (A, **B)** Raw two-colour MINFLUX (2D) images showing the synaptic ribbons labelled with an antibody against CtBP2/RIBEYE (yellow, marked by dotted lines) and the topographic distribution of vesicular compartments marked by the vesicular glutamate transporter VGLUT3 (magenta). (A) and (B) are exemplary images from two different IHCs. Inset in (A) shows confocal overview of the IHC (marked by VGLUT3 labelling) and zoom-in with individual channels showing VGLUT3 (CF660C conjugated secondary antibody) and CtBP2/RIBEYE (Alexa647 conjugated secondary antibody) signal prior to spectral unmixing. Scale bar = 200 nm for MINFLUX images and 5 μm for confocal inset. Inset in (B) shows zoom in of the synaptic ribbon and a schematic illustration of the synaptic ribbon to elucidate possible SV arrangement around the ribbon and the presumable position of the presynaptic membrane (dotted line). **(C)** Molecular counts of VGLUT3 were assigned into clusters using DBSCAN and fitted with circles to depict putative vesicles. **(D)** Diameter of the circular fits of VGLUT3 clusters have been presented in a histogram.

### Nanoclusters of channels and interacting proteins govern AZ and PSD topography

We next assessed the spatial arrangement of the various synaptic proteins to observe if the protein units show some higher-order organisation at the AZ and the PSD. To do so, we used Monte Carlo simulations to compare the cumulative distribution of the nearest neighbour distances, known as G-function, of the measured unit positions with synthetic data of randomly arranged positions confined to the same measured volume and with the same number of units. The simulations were repeated 1000 times for each synapse in order to generate a confidence envelop. A G-function with a curve higher than the simulation envelop indicates clustering behaviour, while one with values below (or to the right of) the simulation envelop, indicates a dispersion pattern. When the measured G-function falls within the simulated envelop for random positions, this means the measured pattern does not deviate from the null hypothesis, which is a random arrangement in our case. With the exception of RIBEYE, all the other synaptic proteins showed clustering behaviour as evident from the positive deviation of their G-functions to the simulated envelops (Fig. 6A, B). In order to derive a binary conclusion from the envelop graphs, we applied the Diggle-Cressie-Loosemore-Ford (DCLF) test (*74*) with probing distances from 10 nm to 18 nm (see Methods). The observed results indicated that in 77.8% of synapses, CaV1.3 units exhibit some degree of clustering (significance level α=0.002; see Fig. 6C and Fig. S6). Data for Piccolino, RBP2 and Homer1 also indicate clustering for 60%, 70% and 90% of synapses respectively, while GluA2 showed the most prominent spatial clustering, having a non-random behaviour for all the imaged synapses. On the other hand, the spatial distribution of RIBEYE units within the synaptic ribbon appeared non-clustered and did not indicate any statistically significant higher-order organisation (Fig. 6B, C; Fig. S6).

**Fig. 6.**
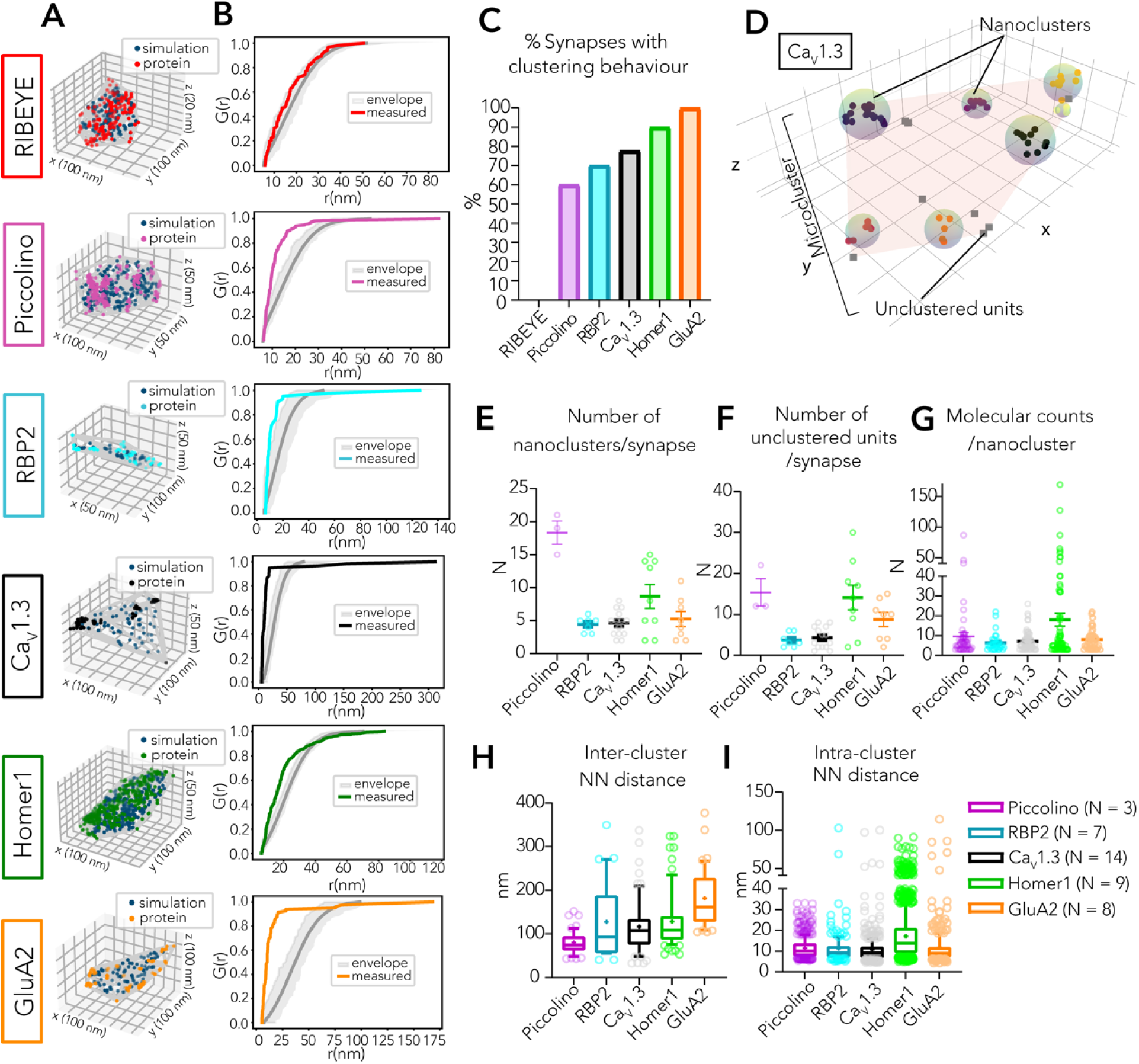
Quantitative analysis of nanoclustering of synaptic proteins. **(A)** Representative 3D projections of measured protein units (coloured) and simulated random positioned units confined to the measured convex hull. **(B)** Respective G-functions of graphs in (A) (coloured) with the envelop representation (shaded in grey) for 1000 Monte Carlo simulations of random distributed units. **(C)** DCLF Test shows clustering behaviour at majority of synapses for all proteins except RIBEYE, suggesting a random distribution of RIBEYE within the synaptic ribbon. **(D)** Plot showing CaV1.3 units after assignment of “nanoclusters” using DBSCAN. Units not assigned into nanoclusters are shown as grey squares; spherical fits representing nanoclusters have radius equal to the distance between nanocluster centroid and the most distant point in the nanocluster measured from that centroid; alpha shape overlay depicts the CaV1.3 “microcluster”; grid boxes are 50 nm along all axes. **(E - I)** Plots depicting mean ± SEM number of nanoclusters per synapse, number of unclustered units per synapse, molecular counts inside each nanocluster, nearest neighbour distance between centroids of individual nanoclusters and nearest neighbour distance between molecular counts within the same nanocluster respectively. For all plots from (E – I), individual data points have been overlaid and represent number of synapses for (E) and (F), total number of nanoclusters for (G) and (H) and total molecular counts from all synapses for (I). Box whisker plots display crosses representing mean values, central band indicating the median, whiskers representing 90/10 percentiles and boxes representing 75/25 percentiles.

To further investigate clustering of Ca_V_1.3, RBP2, GluA2, Homer1 and Piccolino, we implemented DBSCAN on units from synapses for which the DCLF test held true, with minimum number of points to form a cluster = 3 and radius e = mean of nearest neighbour distance of all units/synapse + 2 standard deviations (Fig. 6D). A summary of the data has been presented in Table 2 and Fig. 6E – I. Ca_V_1.3 *microclusters* appeared to be composed of 2 – 8 *nanoclusters* per synapse at an inter- cluster distance (between centroids of nanoclusters) ranging from 32.8 – 381.8 nm (average = 120.6 ± 8.8 nm, S.D. = 71), with an average nanocluster diameter of 47.0 ± 4.2 nm, S.D. = 34.3. Each of these nanoclusters in turn appeared to contain between 3 – 26 channels (average 7 ± 0.6 molecular counts/nanocluster, S.D. = 5) spaced at an average intra-cluster distance of 10.8 ± 0.4 nm (S.D. = 8.2, median = 9.1 nm). On average 1 – 8 individual channels not assigned into a nanocluster were present at each synapse. These estimates closely match with estimates derived from RBP2 which also show 3 – 6 nanoclusters per synapse, each with 3 – 22 RBP2 molecular counts (average 6 ± 0.8 molecular counts/nanocluster, S.D. = 5), spaced at a distance of 11.3 ± 0.6 nm from the nearest neighbour. We found 2 – 11 (average = 5 ± 1) GluA2 nanoclusters/synapse, with on average 9 GluA2 units per synapse not assigned to a nanocluster. GluA2 nanoclusters seem to be composed of 3 – 22 GluA2 units (average = 8 ± 1) at an average distance of 12 nm from the nearest neighbour.

**Table 2.** Summary of quantitative analysis of nanoclusters. Counts (N) = number of synapses for number of nanoclusters/synapse and number of unclustered units/synapse; N = total number of nanoclusters for molecular counts/nanocluster and inter-cluster nearest neighbour distance; and N = total molecular counts from all synapses for intra-cluster nearest neighbour distances.

|  | No. of nanoclusters /synapse | No. of unclustered units /synapse | Molecular counts /nanocluster | Inter-cluster NN distance (nm) | Intra-cluster NN distance (nm) |
| --- | --- | --- | --- | --- | --- |
| **Piccolino** |  |  |  |  |  |
| Mean | 18 | 15 | 10 | 80.5 | 11.4 |
| S.D. | 3 | 6 | 14 | 25.3 | 5.0 |
| SEM | 2 | 3 | 2 | 3.4 | 0.2 |
| Range | 15 – 21 | 12 – 22 | 3 – 87 | 43.6 – 152.0 | 5.5 – 33.0 |
| Median | 19 | 12 | 6 | 74.5 | 10.1 |
| Count | 3 | 3 | 55 | 55 | 530 |
| **RBP2** |  |  |  |  |  |
| Mean | 4 | 4 | 6 | 127.9 | 11.3 |
| S.D. | 1 | 2 | 5 | 83.9 | 9.0 |
| SEM | 0.4 | 0.6 | 0.8 | 15.1 | 0.6 |
| Range | 3 – 6 | 2 – 6 | 3 – 22 | 40.8 – 349.4 | 5.1 – 103.2 |
| Median | 5 | 3 | 5 | 93.0 | 9 |
| Count | 7 | 7 | 31 | 31 | 197 |
| **Cav1.3** |  |  |  |  |  |
| Mean | 5 | 4 | 7 | 120.6 | 10.8 |
| S.D. | 2 | 3 | 5 | 71.0 | 8.2 |
| SEM | 0.5 | 0.7 | 0.6 | 8.8 | 0.4 |
| Range | 2 – 8 | 1 – 8 | 3 – 26 | 32.8 – 381.8 | 4.6 – 100.3 |
| Median | 5 | 4 | 6 | 107.9 | 9.1 |
| Count | 14 | 14 | 65 | 65 | 468 |
| **Homer1** |  |  |  |  |  |
| Mean | 9 | 14 | 18 | 128.4 | 17.2 |
| S.D. | 5 | 9 | 28 | 63.9 | 11.7 |
| SEM | 2 | 3 | 3 | 7.4 | 0.3 |
| Range | 2 – 15 | 2 – 30 | 3 – 169 | 53.4 – 324.1 | 4.9 – 91.1 |
| Median | 8 | 14 | 6 | 108.7 | 13.8 |
| Count | 9 | 9 | 75 | 75 | 1354 |
| **GluA2** |  |  |  |  |  |
| Mean | 5 | 9 | 8 | 182 | 11.7 |
| S.D. | 3 | 5 | 5 | 64.7 | 10.8 |
| SEM | 1 | 2 | 0.8 | 10.0 | 0.6 |
| Range | 2 – 11 | 2 – 15 | 3 – 22 | 104.6 – 377.1 | 4.3 – 114.7 |
| Median | 5 | 10 | 7 | 161.6 | 8.9 |
| Count | 8 | 8 | 42 | 42 | 335 |

### CaV1.3 nanoclusters serve efficient SV release of IHC ribbon synapses

We next employed biophysical modelling to assess if nanocluster-based organisation of CaV1.3 would impart any functional advantages at the IHC ribbon synapse. We simulated Ca^2+^ binding to SV Ca^2+^ sensors (*41*) at the vesicular release site by utilizing the particle-based Monte Carlo method (MCell) (*75*). We assumed that a single fully Ca^2+^-occupied sensor triggers SV release and estimated the release probability by accounting for stochastic influx and diffusion of Ca^2+^ in the presence of buffers estimated for physiological conditions (*18*). First, we modelled a simple scenario where a single SV was coupled to a nanocluster consisting of 12 voltage-gated Ca²⁺ channels. The SV diameter was set to 50 nm (*17*), and the coupling distance (*d*) - defined as the distance between the SV centre and the edge of the nanocluster - was fixed to 15 nm (see Fig. 7A). Six sensors were placed with a polar angle (θ) of 115° empirically as direct experimental estimates of the number and location of the SV Ca^2+^ sensors are currently unavailable. We generated 10 different scenarios of Ca^2+^ channel distribution and simulations were repeated 100 times. The local [Ca^2+^] was estimated with the hit rate per unit area, showing ∼ 40 µM around the sensor index 1 (see supplementary text and Fig. S7). For each sensor, we estimated the effective coupling distance, Rc (see Methods). As shown in Fig. 7B (top panel) when sensor indices are ordered according to the average distance between a sensor and channels, a closer proximity to the nanocluster corresponds to a lower Rc (black line) and a higher number of release events (grey line). The closest sensor (sensor index 1) exhibited an Rc of 23.2 nm, while the other sensors showed values of 32.1 nm, 33.1 nm, 50.6 nm, 51.2 nm, and 59.8 nm, respectively. Most release events were triggered by sensor index 1 (74.5 %), followed by sensor index 3 (14.4 %), sensor index 2 (10.9 %), sensor index 4 (0.2 %), and others did not trigger any release events.

**Fig. 7.**
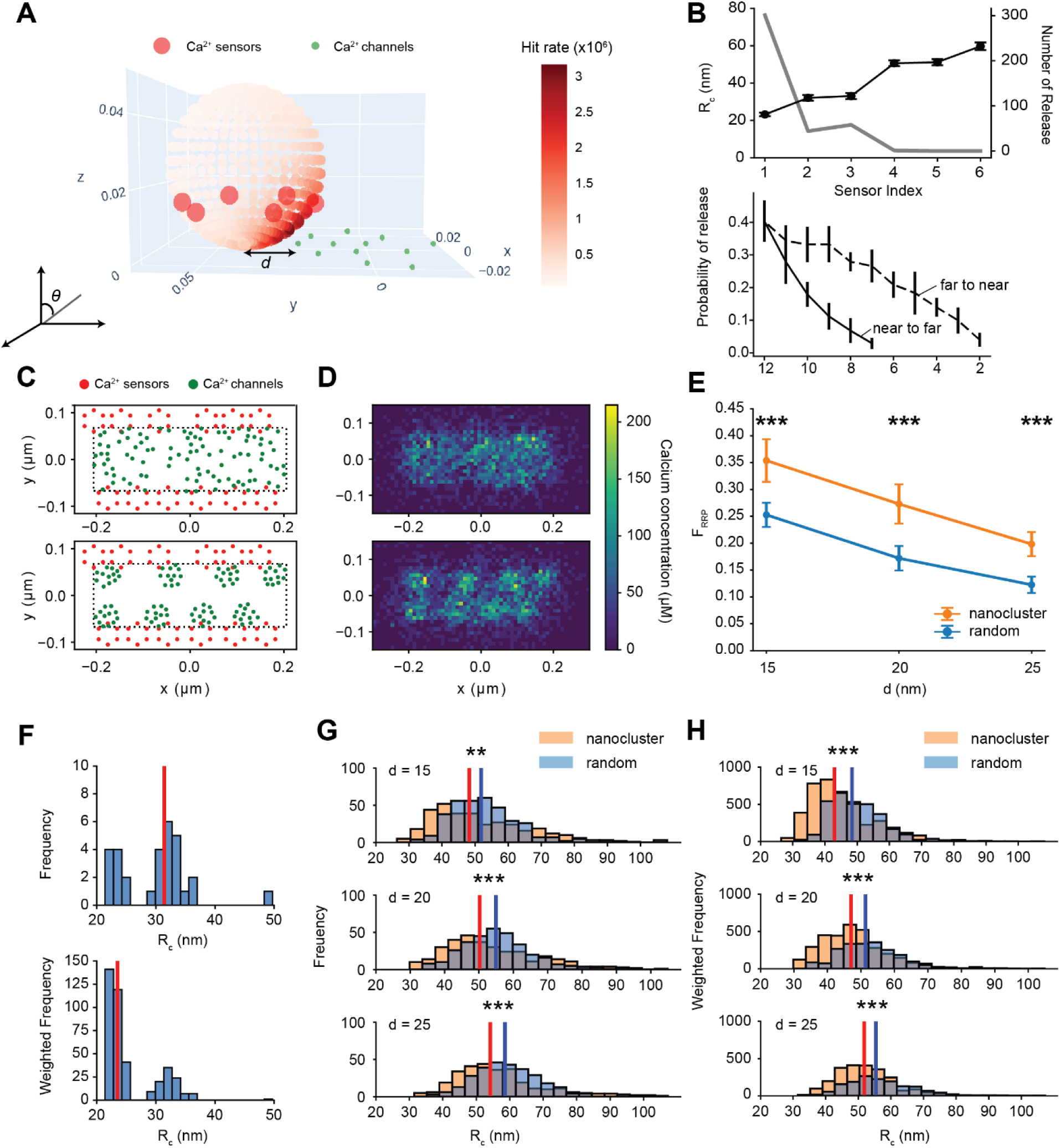
Monte Carlo simulations of Ca^2+^-dependent vesicle release in two Ca^2+^ channel-SV topographies. **(A)** 3D spatial representation of one pair of SV and Ca^2+^ channel nanocluster. Six sensors were placed on the SV surface (red) and detected hit rates per unit area are represented in a colour map on the SV surface. Twelve Ca^2+^ channels are placed into the nanocluster with a diameter of 50 nm (green). **(B)** Top panel shows effective coupling distance, Rc (black) and number of release events for the individual sensors (grey) versus sensor index. Bottom panel shows release probability versus number of Ca^2+^ channels. Ca^2+^ channels were progressively removed based on the distance to sensor index 1. Removing channels closest to furthest from the SV is labelled as “near to far” and from the furthest to the closest as “far to near”. **(C)** Top view of one of 10 different spatial distributions of 96 Ca^2+^ channels in random (top) or nanocluster (bottom) configurations. **(D)** Ca^2+^ concentration ([Ca^2+^]) estimated with a voxel size of 10 nm x 10 nm x 10 nm. Range of z-axis is from 0 nm to 10 nm. Colour bar indicates the range of [Ca^2+^] in µM. **(E)** Release probability as a function of coupling distance *d* for nanocluster (orange) and random (blue) scenarios; error bars indicate standard deviations, and statistical significance is marked (t-test, *** *p < 0.001*). **(F)** Histograms of Rc in single SV-nanocluster case. Red line indicates median value. **(G)** Histograms of Rc at different *d* values (*d* = 15, 20, 25 nm), comparing nanocluster (orange) and random (blue) scenarios; statistical significance is indicated (Mood’s median test, ** *p < 0.01*, *** *p < 0.001*). Red and blue lines indicate median for nanocluster and random scenario respectively. **(G)** Weighted frequency histograms of Rc (incorporating number of release events per bin) for different *d* values, (t- test, *** *p < 0.001*).

**Fig. 8.**
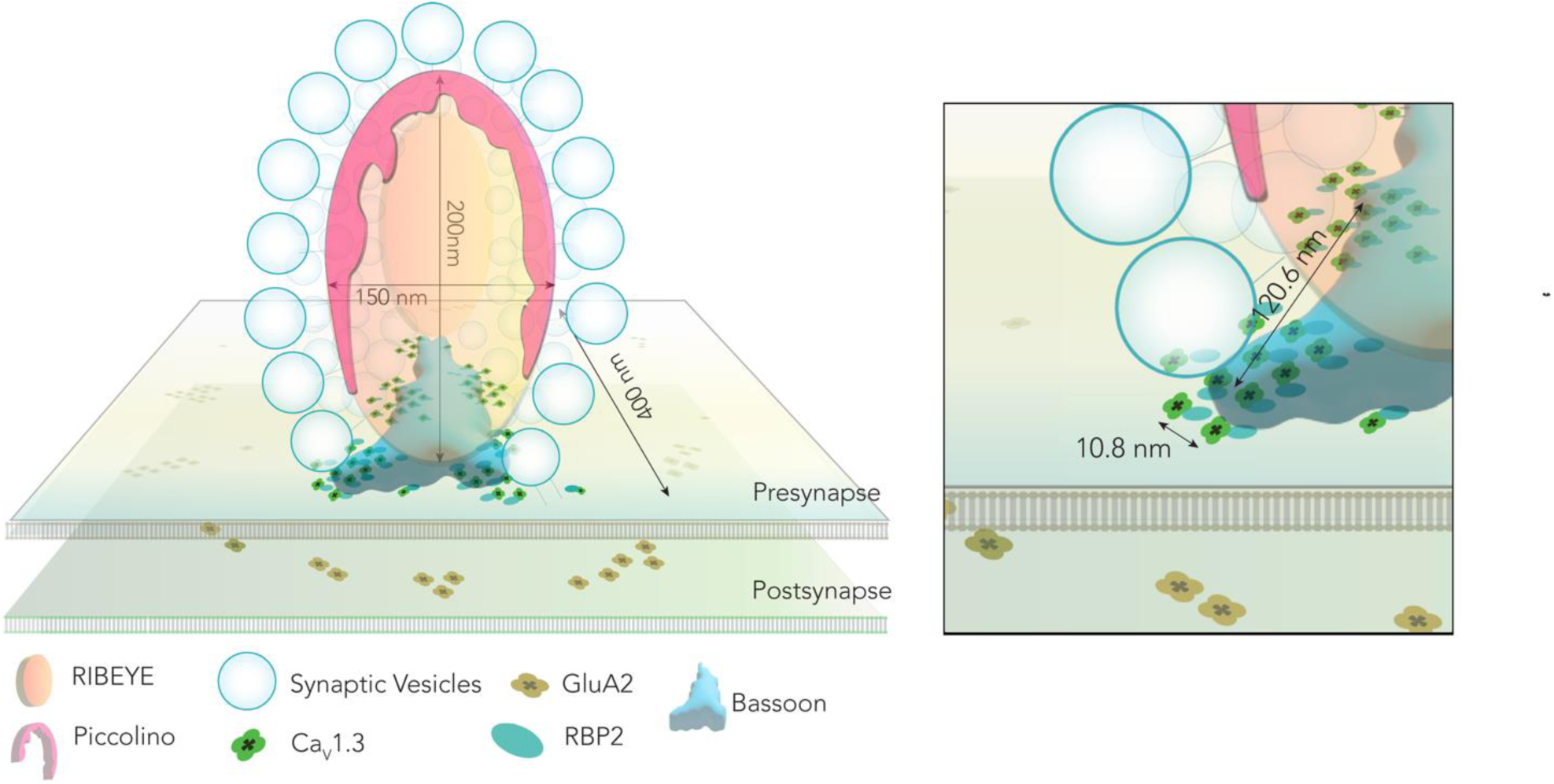
Speculative model of the molecular nanoanatomy of IHC ribbon synapses based on MINFLUX data. Schematic shows the cross-section of the synaptic ribbon along its short axis. CaV1.3 and RBP2 nanoclusters form a stripe-like microcluster underneath the synaptic ribbon. Some individual channels are dispersed in between. RBP2 links CaV1.3 to Bassoon (inferred from published data, as no MINFLUX data were obtained for Bassoon), which localizes underneath the ribbon as well and anchors the ribbon to the presynaptic membrane. A good dozen of release ready SVs sits within 17 nm of CaV1.3 nanoclusters as assumed based on physiology and electron tomography. Two or more SVs may be shared by a CaV1.3 nanocluster and may be positioned towards the outer perimeter of a given CaV1.3 nanocluster or SVs may be assigned to CaV1.3 nanoclusters in a one-to-one stoichiometry. Postsynaptic GluA2 receptors are arranged in a ring- like topography engulfing the presynapse for efficient glutamate detection.

As shown in Fig. 7B (bottom panel), the release probability within 2 ms was 0.4 ± 0.06. Notably, progressive removal of Ca²⁺ channels, starting with those closest to the sensor index 1, led to a rapid decline in release probability (Fig. 7B, bottom panel). The removal of the two closest Ca²⁺ channels, “near to far”, reduced release probability by 55 % (from 0.4 to 0.18), and the removal of five Ca²⁺ channels resulted in nearly zero release. Interestingly, progressive removal of Ca^2+^ channels furthest from the sensor index 1, “far to near”, led to slower decline in release probability. In such a scenario, deleting 7 Ca²⁺ channels resulted in a comparable decrease in the release probability as deleting 2 Ca²⁺ channels in the “near to far” condition. Furthermore, having only 2 closest Ca²⁺ channels did not trigger any release events. This may indicate that in addition to the 2 – 3 “private” channels (*21*) closest to the sensor which are the primary drivers of release, surplus channels in a nanocluster may also contribute e.g. by building a high microdomain [Ca^2+^] and thereby, depleting buffer molecules, and likely promoting Ca^2+^ dependent SV replenishment by “private” channels (*76*).

Next, we extended our simulations to physiologically relevant scenarios with multiple nanoclusters taking advantage of physiological channel counts (*18*) and morphological evidence for channel topography (this study, (*18*)). To evaluate the functional efficiency of nanocluster organization, we estimated the fraction of the readily release pool (RRP) released within 2 ms in two topographical scenarios: i) random Ca^2+^ channel distribution, and ii) nanoclusters. In both scenarios, we distributed 96 Ca_V_1.3 Ca^2+^ channels within a rectangular active zone with dimensions 410 nm × 135 nm. The minimum inter-channel distance was set to 11 nm. For the nanocluster scenario, 12 channels were distributed within a 50 nm diameter and 8 nanoclusters were distributed as two lines, where each nanocluster was placed at the edge of the AZ. The nanocluster centre was fixed at 42.5 nm and -42.5nm along the y-axis, while nanoclusters were randomly positioned along the x-axis. We positioned 12 docked SVs around Ca^2+^ channel clusters (*17*) (Fig. 7C, D). For each scenario, we further generated 10 possible configurations with different channel distributions and SV arrangements. Each configuration for a scenario was simulated 100 times (i.e., 1000 iteration for nanocluster arrangement with a *d* of 15 nm). We tested three coupling distances (*d*): 15 nm, 20 nm, and 25 nm. Other model parameters including buffer concentrations, Ca^2+^ channel kinetics, and Ca^2+^ binding kinetics of the Ca^2+^ sensor on the SV were based on previous physiological estimates (see methods, Table S6). To investigate the effect of spatial coupling between the SV Ca^2+^ sensors and Ca^2+^ channels on synaptic efficacy, we used the fraction of released RRP (*FRRP* within τ of RRP release) as a proxy of the release probability. The physiological estimates of *FRRP* at 2ms range from 0.09 - 0.27 (see methods and (*77*)). Here, we found the *FRRP* for random arrangement of channels to be 0.25 ± 0.02, 0.17 ± 0.02, 0.12 ± 0.02 (for *d* = 15, 20 and 25 nm respectively). As expected, *FRRP* decreases with increasing coupling distance *d*. For the nanocluster scenarios, the predicted *FRRP* were 0.35 ± 0.04, 0.27 ± 0.04, 0.2 ± 0.02 at *d* = 15, 20, and 25nm, respectively. For a given coupling distance, our simulations predict that a nanocluster arrangement of Ca^2+^ channels provides more efficient SV release when compared with randomly distributed channels (\*\*\**P* < 0.001, t-test), (Fig. 7E).

To further investigate the relationship between Rc and release probability, we plotted histograms of Rc first for the single SV-nanocluster scenario, where only the coupling distance values associated with release events were included (Fig. 7F). This shows that SV release events were triggered when the coupling distance was within 50 nm (median of 31.4 nm). To better reflect the relation between the Rc and the number of release events, we additionally plotted a weighted histogram taking into account the number of release events (weighted median of 23.5 nm). This revealed two distinct groups: one with shorter Rc around the weighted median from sensor index 1, corresponding to the majority of release events (74.5%), and another with relatively higher Rc from sensor indices 2 and 3 (25.3%), corresponding to the remaining release events.

We next analysed histograms of Rc for the simulated nanocluster versus random arrangement of channels at the AZ (Fig 7G). The nanocluster scenario exhibited shorter Rc than the random distribution; for instance for coupling distance of 25 nm, median is 59 nm for random *versus* 54 nm for nanoclusters (\*\*\**P* < 0.001, Mood’s median test). Additionally, generating weighted histograms accounting for the number of release events per bin (Fig 7H) revealed that shorter Rc corresponded to higher release probabilities; median of 55 nm in random *versus* 51 nm for nanoclusters (\*\*\**P* < 0.001, t-test). This highlights the functional advantage of a nanocluster-based organisation and that in physiology a difference of 4 – 5 nm in Rc might significantly affect release dynamics.

## Discussion

In this study, we employed 3D MINFLUX to elucidate the molecular nanoanatomy of IHC ribbon synapses. We established immunolabeled semithin Epon-embedded cochlear sections which provide minimal background and allow immobilization of synapses on the coverslip for efficient single molecule localisations with MINFLUX. 3D MINFLUX revealed the densely packed, non- clustered arrangement of RIBEYE molecules inside the synaptic ribbon, with Piccolino distributed along the membrane-distal apex of the ribbon, enveloping it in the form of an arch. Using two- colour MINFLUX imaging of VGLUT3 and synaptic ribbons, we visualize putative SVs and other VGLUT3-positive organelles/membranes around the ribbon which, thus far, were only accessible to EM. Our data revealed nanoclusters of Ca_V_1.3 voltage-gated Ca^2+^ channels and the multi-domain presynaptic AZ protein RBP2 that were typically arranged in a narrow stripe along the long axis and underneath the ribbon. Nanoclusters of Ca_V_1.3 seem to be composed of 3 – 26 channel units, interspersed at an average distance of 10.8 nm from the nearest neighbouring channel.

Mathematical modelling indicates that such a nanocluster-based organisation of CaV1.3 enables a higher release probability by shorter effective coupling distance, contributing to the efficiency of synaptic sound encoding in the ear. Our MINFLUX data also sheds light on the topography of postsynaptic Homer1 and AMPA receptors, with hundreds of Homer1 units arranged in densely packed, flat patches at the postsynaptic density and nanoclusters of GluA2 receptor units assembled in a ring, presumably at the periphery of the postsynaptic density.

### Molecular nanoanatomy of the IHC ribbon synapse

Our MINFLUX study supports and extends previous structural studies of the IHC ribbon synapses using STED imaging and/or EM. Our data agree with morphology and dimensions of microclusters of CaV1.3 channels (*6*, *11*, *14*, *18*, *20*), RBP2 (*20*), Piccolino (*11*, *54*), RIBEYE (*3*, *21*, *54*, *66*), and GluA2 (*65–67*). However, while RIBEYE has been previously proposed to assemble in a staggered array inside the ribbon scaffold (*8*) and, occasionally, lamellar substructure has been observed in mature synaptic ribbons with EM (*3*, *52*, *53*), we did not observe any such lamellar or staggered organisation of RIBEYE assembly inside ribbons of mature mice. Instead, we found RIBEYE to show a distribution statistically indistinguishable from spatial randomness inside the ribbon. We cannot exclude this finding to be specific to IHC ribbons studied here. Moreover, future cryogenic electron tomography and subtomogram averaging might help further addressing the molecular structure of this enigmatic nanomachine.

Previous observations from MINFLUX imaging of the ribbon synapses in retinal photoreceptors (*33*), indicated an arrangement of AZ proteins Bassoon, RIM2, CaV1.4 Ca^2+^ channels and ubMunc13-2 in two distinct parallel lines along each site of the rod ribbon. This has been taken to suggest a regularly ordered repetition of a molecular and structural correlate of SV release sites in linear tracks on the two sides of the synaptic ribbon. This seemed reminiscent of the classic “ribs and pegs” model of the AZ from EM studies at the frog neuromuscular junction (*30*, *78*). In contrast, MINFLUX imaging of RBP2, CaV1.3 and even RIM2 at IHC AZ show a nanocluster topography typically forming single stripe-like microclusters. MINFLUX imaging also indicated more complex microclusters which can appear linear upon rotation. We note the possibility that two proximal stripes could not be parsed out due to sparse localizations and the small size of the IHC AZ compared to rod photoreceptor AZ. However, what seems more likely is that the occurrence of double stripes is simply rare for IHC AZs and majority of the AZs actually contain just one narrow stripe (at up to 60% of synapses) as also supported by numerous observations from STED microscopy (*6*, *11*, *18*, *20*). Indeed, we observed two defined short stripes of CaV1.3 localizations at one synapse (Fig. 2K), whereas other complex shaped clusters could also subjectively be considered as non-parallel lines (e.g. Fig S5A). This appears in agreement with occasional findings of two stripe-like/fat line shaped CaV1.3 microclusters by STED (*18*) and is also consistent with finding multiple stripes of intramembrane particles in freeze-fracture EM images of the larger frog saccular hair cell active zone (*36*).

### Mapping SVs using light microscopy

The molecular machinery underlying IHC exocytosis is unconventional (*1*): the role of canonical neuronal SNAREs is controversial (*79*, *80*); Synaptotagmin 1 (*81*, *82*), Complexins (*83*), Munc13s (*16*) and Munc18s (*84*), Synaptophysin 1 and Synapsin 1, 2 (*85*, *86*), VGLUT1 and VGLUT2 (*68*, *69*) seem to be absent. Instead, IHC synapses rely on VGLUT3 (*68*, *69*) and on Otoferlin that is involved in SV tethering, docking, Ca^2+^ sensing for fusion and exo-endocytosis coupling (review in *89*, *90*) and defective in hereditary deafness DFNB9 (*89*). Otoferlin distributes broadly in IHCs (*42*, *71*, *72*) and, aside from SV cycling, seems also involved in constitutive membrane trafficking (*72*). The Otoferlin labelling we achieved in the semithin section thus far did not yet lead to efficient MINFLUX imaging. Further optimisation of stainings or the use of mouse lines expressing tagged- Otoferlin will serve future MINFLUX studies. Here, we embarked on detecting SVs by labelling for VGLUT3, which is a bona fide SV marker in IHCs (*68*, *69*, *71*, *72*, *84*). Our two-colour MINFLUX data shows the expected broad distribution of VGLUT3 in the synaptic IHC pole and nicely depicts putative SV localizations around the synaptic ribbon and near the membrane. Further validation by co-labelling for other SV markers like Otoferlin will help making MINFLUX available as an approach to study SV distributions at the AZ including at different states of activity, complementing electron tomographic studies (*13*, *17*) with molecular information. This will be particularly meaningful for assessing the spatial relationship of SV release sites, SV Ca^2+^ sensors and the CaV1.3 channels triggering SV release.

### Determining “molecular counts” of synaptic components using MINFLUX

The first estimates of Ca^2+^ channels at IHC ribbon synapses came from a study in the frog which used non-stationary fluctuation analysis on whole-cell Ca^2+^ tail currents and reported on average 80 Ca^2+^ channels per AZ (*36*). A subsequent study in mouse IHCs also reported a similar estimate of 80 Ca^2+^ channels per AZ using fluctuation analysis (*35*). A later study employed imaging of presynaptic Ca^2+^ influx in combination with patch-clamp and performed (i) optical fluctuation analysis and (ii) correlation of Ca^2+^ influx at a single AZ and whole-cell Ca^2+^ influx while locally depleting free extracellular Ca^2+^ via iontophoresis at that single AZ using the Ca^2+^ chelator EGTA (*18*). Through both approaches, a wide range of channels/AZ was reported: 20 – 200 channels/AZ (median of 60) for the first and 30 – 300 channels/AZ (median of 120) for the latter. Quantitative analysis of our data suggests between 9 – 70 (mean = 35 ± 4; median = 34) molecular counts of CaV1.3 per synapse in IHCs. While our estimates lie within range of previously published functional estimates in IHCs (closer to the estimates predicted by optical fluctuation analysis), they likely underestimate the number of CaV1.3 per synapse probably due to limiting labelling efficiency. Molecular counts and topography of RBP2 match well with those of CaV1.3, consistent with the proposed role of RBP2 in positively regulating CaV1.3 abundance at the IHC ribbon synapse (*20*, *90*). Studying the relative topographies of CaV1.3, RBP2, and Bassoon remains an important objective for future multicolour MINFLUX studies. It is worth noting that the number of CaV1.3 channels, and the resultant size of CaV1.3 channel microclusters (or intensity of Ca^2+^ influx in the case of functional Ca^2+^ imaging studies), scales with size of the synaptic ribbon (*11*, *18*, *91*, *92*). This becomes even more relevant since IHCs display a remarkable position-dependent heterogeneity in terms of synapse size and properties along with the firing properties of corresponding SGNs (for review see *100*).

Another limitation of the current study is the linkage error of ∼20 nm when localizing proteins by indirect immunolabeling (for review see *101*). Even though emitter localisations were performed with high optical precision, the linkage error compromises the accuracy of determining protein localisations themselves. Unfortunately, nanobodies, that could alleviate this problem, were largely unavailable for the targets of our study, or generated weak labelling not supporting MINFLUX imaging. Moreover, our attempts using secondary nanobodies in combination with primary antibodies also resulted in weak labelling which did not result in successful MINFLUX imaging. Future work employing mice expressing tagged proteins such as CaV1.3 with a HaloTag (*95*) will help to determine localisations and molecular counts more accurately.

### Nanotopography of Ca^2+^ channels using MINFLUX – what can we learn?

Our 3D MINFLUX data reveals that the CaV1.3 channels and RBP2 appear as discrete nanoclusters positioned along the length of the synaptic ribbon. While we cannot entirely rule out the possibility of clustering due to fixation, a recent study shows that the modulatory C-terminal domain of CaV1.3 is sufficient to promote nanoscale channel clustering even in live iPSC-derived cardiomyocytes (*96*). Moreover, the spatial arrangement of RIBEYE units inside the ribbons as seen in our MINFLUX data does not show such a nanoclustering also speaks against this possibility. Our data converge with an average of 7 CaV1.3 units per nanocluster, at a mean distance of 10.8 nm from the nearest neighbouring channel within the same nanocluster similar to the nanoclusters reported (*96*). Moreover, a report from CNS synapses using EM on SDS-digested freeze-fracture replica labels also arrived at an average estimate of ∼9 CaV2.1 channels per nanocluster, with each nanocluster tightly coupled in a one-to-one stoichiometry with a vesicular release site (*28*). IHCs are predicted to have 10 – 15 release sites per AZ, with approximately 80 – 120 channels per AZ (*14*, *18*, *71*, *97*), which would give a similar figure of 8 channels per release site, of which 1-3 “private” CaV1.3 channels exert a Ca^2+^ nanodomain-like control of release with an effective coupling distance of ∼17 nm (*21*, *37*). However, our data reports an average of 5 nanoclusters/synapse (range = 2 – 8) with additional individual/pairs of CaV1.3 units that we did not assign into a nanocluster (between 1 and 8 per synapse). Therefore, we refine our prior suggestion of SV release sites lining the circumference of the CaV1.3 microcluster with an effective coupling distance of the of ∼17 nm from “private” CaV1.3 channels in the periphery of the cluster (*21*, *37*) and propose SV release sites to be positioned around the circumference of CaV1.3 nanoclusters. In such a scenario, we envision that a subset of channels within the nanocluster would be within the described effective coupling distance to the Ca^2+^ sensor of SV fusion. Such a perimeter model has also been recently proposed for the mature inhibitory basket cell-Purkinje cell synapse of the cerebellum, which also show a tight Ca^2+^ nanodomain control of exocytosis (*27*). The CaV1.3 nanoclusters and associated release sites at the IHC synapses represent modular implementation of maximal synaptic strength which offers versatility. IHCs likely not only vary in the number of these modules but this way also might tune the coupling of Ca^2+^ influx with SV release. Indeed, paired pre- and postsynaptic patch-clamp recordings and combined imaging of presynaptic Ca^2+^ signals and glutamate release demonstrate variations of the prevailing Ca^2+^ nanodomain coupling across AZs (*38*, *39*, *77*).

It would also be interesting to look at CaV1.3 nanoclustering prior to the onset of hearing, when Ca^2+^ microdomain-like control of exocytosis is prevalent (*21*), or using genetic models with perturbed RIBEYE, Bassoon or Piccolino which show disrupted Ca^2+^ channel clusters and/or diffuse Ca^2+^ influx (*6*, *11*, *14*, *18*). Our data argues against the possibility of an exclusion zone (*34*) at the mature IHC ribbon synapse. Future experiments with multicolour MINFLUX to directly visualize CaV1.3 channel topography with respect to vesicular release sites/vesicular markers would add more clarity. However, this might be challenging as two-colour MINFLUX becomes difficult when the abundance of fluorophores is considerably disproportionate for two different proteins and blinking events from one would merely dominate over the other (e.g.: VGLUT3 over CaV1.3). Future studies combining DNA-PAINT (DNA points accumulation for imaging in nanoscale topography) and MINFLUX (*98*) could be used to perform multiplexing to investigate multiple synaptic structures in the same sample using MINFLUX.

Lastly, at CNS synapses, presynaptic AZ nanoclusters have been shown to be positioned in tandem with postsynaptic AMPA receptor clusters generating synaptic nanocolumns (*23*). We did not exclusively probe the existence of such transsynaptic nanocolumns at the IHC ribbon synapse using 2-colour MINFLUX. However, it seems likely that the IHC-SGN synapse has a different implementation where a presynaptic AZ extending to just about 400 nm along its length is engulfed by a “ring-like detector” of postsynaptic AMPA receptor clusters which extend to even ∼900 nm. This way, the distribution of AMPA receptor nanoclusters might maximize efficiency for detection of glutamate as proposed before (*65*, *67*).

## Materials and Methods

### Tissue processing and sectioning

All animal experiments complied with guidelines from the University of Göttingen Board for Animal Welfare and the Animal Welfare Office of the State of Lower Saxony. Cochleae from 2- week-old (postnatal day P14 – P21) C57BL6/J mice were dissected in ice-cold phosphate buffer saline (PBS) and immediately fixed in 4% formalin (in PBS) on ice for 40 – 60 minutes. For samples fixed with glyoxal (*51*), fixation solution (3% w/v) was prepared as follows (4 ml solution mix): 2.835 ml ddH2O, 0.789 ml ethanol, 0.313 ml glyoxal (40% stock solution from Sigma-Aldrich),

0.03 ml acetic acid; solution was vortexed and brought to pH 4/5 by adding 1 M NaOH. Samples were then fixed for 30 min on ice and then 30 min at room temperature. Following fixation with either protocol, samples were washed thoroughly in PBS three times (5 – 10 minutes each wash) and then decalcified using Morse’s solution (10% sodium citrate + 22.5% formic acid) for 20 – 30 min. Cochleae were washed again for three times (5 minutes each) in PBS.

After decalcification, the cochleae were embedded in Epon epoxy resin after dehydration using acetone and polymerized for 24 hours at 60°C. 1 μm sections were cut using Diatome Histo diamond knife and a Leica UC7 microtome. About 3 – 4 such sections were placed on 18 – 24 mm diameter coverslips (thickness = 1.5H i.e., 170 ± 5 μm). The coverslips containing the sections were heated on a hot plate (60°C) for about 15 – 20 min to immobilize the sections on to them. These were then used for immunostainings or could be stored at room temperature until needed.

### Immunohistochemistry

Prior to immunostaining, we removed the Epon resin by treating coverslips containing sections with a saturated solution of NaOH in absolute ethanol (*50*) for 15 min at RT. This was followed by thorough washing with PBS (3 – 4 times). Blocking and permeabilisation was performed in a humidified wet chamber with GSDB (goat serum dilution buffer: 16% normal goat serum, 450 mM NaCl, 0.3% Triton X100, 20 mM phosphate buffer, pH∼7.4) for 1 h at room temperature. We then incubated the coverslips with primary antibodies (diluted in GSDB, refer to Table S1) overnight at 4°C. Afterwards, the samples were washed three times for 10 min in a wash buffer (450 mM NaCl, 0.3% Triton X 100, 20 mM phosphate buffer, pH∼7.4). We then incubated the coverslips with respective secondary antibodies (diluted in GSDB, refer to Table S2) for 1 hour in a light-protected humidified chamber at room temperature. Finally, we washed the samples three times for 10 min in wash buffer and two more times with PBS. Samples could be left in PBS at 4 C° for around one-two weeks until MINFLUX imaging.

### MINFLUX imaging

On the day of MINFLUX imaging, the coverslips were washed in PBS and incubated for 5 minutes with 150 nm gold nanoparticles (BBI solutions, EM.GC150), which serve as fiducials during MINFLUX image acquisition to enable sample stabilization and beamline monitoring. Excess of the gold nanoparticles were removed by washing the samples in PBS three times. We then added 10mM MgCl2 for 5 minutes on to the coverslips which enhances adhesion of gold nanoparticles to the coverslip and the sample, followed by three washes in PBS. Samples were mounted in freshly prepared GLOX buffer [50mM Tris-HCl, 10mM NaCl, 10% (w/v) glucose, catalase (64 μg/ml), glucose oxidase (0.4 mg/ml) and 10-25 mM mercaptoethylamine (MEA, pH = 8.0)] for imaging and the coverslips were finally sealed using twinsil (picodent). Imaging was performed on an Abberior 3D MINFLUX system (Abberior Instruments GmbH), as described before (*99*). Briefly, the microscope is built on a motorized inverted microscope body (IX83, Olympus) with CoolLED illumination for epifluorescent imaging via the eye piece. The system is equipped with 405 nm, 485 nm, 561 nm, and 640 nm laser lines, as well as a 980 nm IR laser for the active stabilisation system. Images were acquired with a 60x 1.42 NA UPLXAPO or a 100X 1.45 NA UPLXAPO oil objective (Olympus). Detection was performed with two APDs in spectral windows Cy5 near (650 – 685 nm) and Cy5 far (685 – 720 nm), which allows for simultaneous two colour imaging. Sites of interest were identified with a 488 co-stain via the eyepiece, with ribbons located confocally either directly from the 647 stain or via a 546 co-stain. Regions of interest were selected for MINFLUX measurement and acquisition proceeded with the standard MINFLUX 2D and 3D imaging sequences. Key sequence parameters are listed in Tables S3 and S4 for 2D and 3D acquisitions respectively. UV activation was applied as appropriate to maintain single molecule blinking behaviour.

### MINFLUX data analysis and statistics

Data (drift corrected using beamline monitoring) were imported from *iMSPECTOR* (Abberior Instruments) as .npy files into *pyMINFLUX* (*100*) where we filtered for quality parameters Effective Frequency at Offset (EFO, rejecting > ∼70000) and Centre Frequency Ratio (CFR, rejecting > ∼0.65). MINFLUX images displayed were created using ParaView software 5.11.2 (*101*) after filtering for EFO and CFR filtering on *pyMINFLUX*. Dimensions of synaptic structures reported were measured using the *ruler* function on ParaView. Confocal images were processed using ImageJ and were adjusted for brightness and contrast. Numerical data was sorted using MS Excel and graphs were created using GraphPad Prism or IGOR Pro 7 (Wavemetrics). Figures were compiled using Adobe Illustrator.

Spectral separation of the two colours was performed based on DCR (Detector Channel Ratio) values:

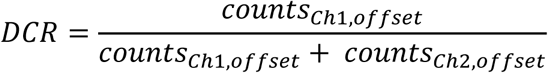

The DCR values were weighted by the corresponding ECO (Effective Counts at Offset) values for each iteration beginning from the headstart. This way, we accounted for any inhomogeneity in the background signal. Then, a two-component Gaussian Mixture Model was fitted to the resulting DCR values and the assigned probability of belonging to distribution 1 or 2 was retrieved. This allowed us to assign a localisation to one colour if its probability was greater than 95%, otherwise the localisation was assigned to a ‘not defined’ category. Afterwards, all the localisations belonging to the same trace, i.e. with the same TID, were assigned to the category corresponding to the majority of the localisations in that trace. In this way all the localisation in one trace would correspond to a single colour. Traces for which majority of the localisations were not defined were not used for visualising and analysing the two-colour MINFLUX data.

The localization precision along each dimension was also determined after pre-processing of data using *pyMINFLUX* and has been reported as the median ± standard deviation of deviations derived from all localizations per Trace ID (TID). The localization precision was typically about 4 to 6 nm along all dimensions. Data from MINFLUX images with more than one synaptic structure were separated into individual data files for each synapse by sorting the coordinates on MS Excel. Filtered data were exported from *pyMINFLUX* as .csv files and were ultimately analysed using a custom written *Python* script. We visualized the raw data using Plotly Express (Plotly Technologies Inc. Collaborative data science, Montréal, QC, 2015. https://plot.ly.) and discarded TIDs with ≤ 4 localizations. Next, we determined the centroid of each TID by averaging coordinates of all localizations per TID and then performed DBSCAN (Density Based Spatial Clustering of Applications with Noise, (*102*)) using a package from *scikit-learn* (*103*) with the minimum number of points to form a cluster = 1. The radius ε of the cluster was calculated as follows (*57*):

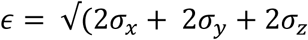

The ranges of ε were as follows (in nm): 5.5 – 5.6 for RIBEYE, 5.8 – 6.1 for Piccolino, 4.9 – 6.0 for RBP2, 4.4 – 6.4 for CaV1.3, 4.9 – 5.8 for Homer1 and 2.9 – 5.7 for GluA2. After removing centroids not pertaining to the synaptic structure and originating from background localizations, we determined the molecular counts (“units”) per synaptic structure and calculated the shortest neighbour distances between them. We used *alphashape 1.3.1* toolbox to fit alpha shapes to the coordinates of units and determined volume estimates of the synaptic structures. The alpha parameter was optimized for complete fitting to structure of interest and was kept consistent for every protein: 0.01 for RIBEYE and Piccolino, 0.001 for RBP2, CaV1.3 and GluA2 and 0.003 for Homer1. For cluster analysis of putative vesicles, DBSCAN was implemented on VGLUT3 units with minimum number of points = 6 and ε = 20. Circle fits have radii equal to the Euclidean distance between the centroid of the individual VGLUT3 clusters and the most distant point in the cluster.

In order to assess the 3D spatial arrangement of the protein units, we performed Monte Carlo simulations of randomly distributed units confined to the convex hull of the measured protein units for each synapse. We applied the constraint that the random units could not be closer to each other with a distance smaller than the calculated localization precision. The number of random units was also constrained to be same as the number of measured units. In this way, we adapted our null hypothesis to a more realistic scenario than the Complete Spatial Randomness, considering the dimension of the measured proteins, their number and the estimated volume that comprise them. The simulations were repeated 1000 times for each synapse and their spatial behaviour was described using the cumulative distribution of the nearest neighbour distances, also known as the G-function. To extract a binary result out of the G-function graphs, i.e., random or non-random, we applied a custom *Python* implementation of the DCLF envelop test (*74*) for distances in an interval ranging from 10 – 18 nm. For analysis of nanoclusters, we further implemented DBSCAN on data containing protein units with the following parameters: minimum number of points to form a cluster = 3 and radius ε = mean of nearest neighbour distance of all units/synapse + 2 standard deviations. Inter-cluster distances reported were calculated as the nearest neighbour distances between centroids of each nanocluster.

### Biophysical Modelling of CaV1.3 nanoclusters

#### Estimation of the fraction of released RRP

Based on the depletion time constant of RRP of high and low spontaneous release (SR) fibres from a previous study (*77*), we estimated the fraction of RRP released within 2ms (*P*_*RRP*_) using the following power-law equation. Assuming that the RRP depletion follows a power-law,

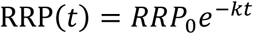

where *RRP*_0_ is the initial number of RRP, *t* represents time, and *k* is the rate constant defined as the inverse of time constant 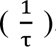 Using this model, the fraction of RRP released within 2 ms was calculated as

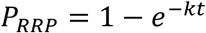

For high SR, the depletion time constant (τ) was estimated to be 6.347 ± 1.096 ms and for low SR, the time constant was 20.88 ± 5.063 ms (*77*). Accordingly, the range of *F*_*RRP*_ at 2ms is from 0.09 (low SR) - 0.27 (high SR).

### Stochastic simulations of Ca^2+^ reaction-diffusion and SV Release

We implemented stochastic simulations of Ca^2+^ reaction-diffusion dynamics and SV release using MCell 3.4 (*75*). A presynaptic active zone (AZ) model was constructed based on a rectangular stripe-shaped Ca^2+^ channel cluster (410 nm × 135 nm) with 96 Ca^2+^ channels and 12 synaptic vesicles. Two coupling scenarios, random distribution and nano cluster distribution, were modelled to investigate spatial dependencies in SV release. Ca^2+^ channel gating was modelled with a three state Markov chain with a parameter set from the previous IHC AZ experimental data (*18*). For Ca^2+^ triggered vesicle fusion kinetics, a seven state Markov chain model was used with a five Ca^2+^ binding steps followed by a vesicle fusion (*41*). The refilling rate was additionally set based on the experimental data, showing the sustained release rate (*71*). The space of the simulated AZ was defined as a hemisphere with a diameter of 1 μ*m* considering that the mean of the nearest neighbour distance of synapses is around 2 μ*m* (*66*). Within the space, fixed and mobile buffers were included (see Table S6). The fixed buffer concentration was set to 610 µM and ATP was included with a concentration of 68 µM. EGTA and BAPTA were set to 800 µM and 0.4 µM, respectively to reflect Ca^2+^ buffering condition in IHCs with both fast and slow binding kinetics (*18*). The bottom of the space was reflective to molecules and the spherical boundary of the space was absorptive for Ca^2+^ ions and we clamped the concentration of mobile buffers BAPTA and EGTA at this surface in order to emulate the refilling of buffers upon Ca^2+^ influx. The clamped concentration was set for both unbound and bound buffers where bound concentration was calculated by multiplying the buffer concentration with bound fraction ([Ca^2+^]·Kon / (Koff + [Ca^2+^]·Kon)) and unbound fraction was calculated as 1 – bound fraction. The single Ca^2+^ current was 0.137 pA (*35*, *104*) with the open channel probability of 0.4, which is the half of the experimental data, considering the absence of BayK8644 in the current simulation condition (*18*, *35*). Each coupling condition was simulated with 10 different scenarios and each scenario was repeated 100 times (total of 1000 simulation iterations for each scenario). Each simulation was with a duration of 2 ms with a time step of 10 ns to match the average diffusion distance of Ca^2+^ at every time step is within nanoscale (∼ 3.37 nm computed via 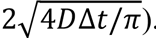). For each iteration, we counted each SV release event when one of SV Ca^2+^ sensors triggered release.

### Ca^2+^ channel gating

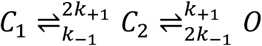

where C1, C2 represent closed channel states, and O represents open channel state. The gating rates k+1 and k-1were set to 1.78 ms^-1^ and 1.37 ms^-1^, respectively, assuming the membrane potential Vm = - 17 mV and the open probability is 0.4 (*21*).

### A kinetic model of five Ca^2+^ sensor binding

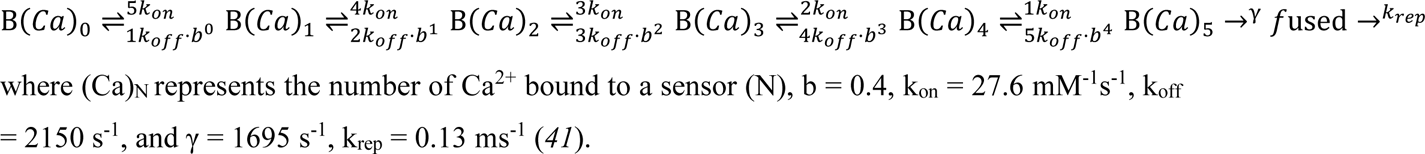

### Estimation of R_c_

To estimate R_c_, we weighted a distance between a sensor and a channel with a Ca^2+^ binding ratio as follows (*37*).

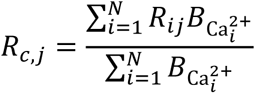

where *R*_*c,j*_ is the effective coupling distance of a sensor *j*, *R*_*ij*_ is the distance between a sensor *j* and a Ca²⁺ channel *i, B*_*Ca*^2+^_i__ is the number of Ca²⁺ binding induced by Ca²⁺ channels *i* to a sensor *j*, *N* is the number of channels.

### Statistical Testing

All statistical analyses to test significant difference in *F*_*RRP*_ were performed using Python packages. We first assessed the normality of each dataset using the Shapiro–Wilk test, given the small sample size and the results indicated that each dataset followed a normal distribution. For comparisons between the random and nanocluster conditions at the same *d* value, we performed a two-sample t-test with equal variance, which was tested with Levene’s test. For comparisons between three *d* values within the same condition, we first conducted a one-way analysis of variance (ANOVA), which was followed by pairwise t-tests. For multiple comparisons, we applied the Bonferroni correction. For comparing the median of the effective coupling distance between random and nanocluster, we used a Mood’s median test. For comparing the weighted median of the effective coupling, we first performed a bootstrap resampling (N=100) by weighting with the number of release events. Then, we compared effective coupling distances between two groups with a two-sample t-test on the distributions of bootstrapped weighted medians.

## Acknowledgments

We would like to thank Dr. Isabelle Jansen and Dr. Ulf Matti (Abberior GmbH) for initial contributions during the project and Dr. Constantin Pape for feedback regarding analysis scripts. We are grateful to Sandra Gerke and Christiane Senger-Freitag for expert technical assistance and Patricia Räke-Kügler for the administrative support during this study. We would also like to thank Prof. Erwin Neher and Prof. Silvio Rizzoli for their feedback and to Dr. Jakob Neef for support throughout the course of this study. We would like to thank Prof. Annalisa Scimemi for providing feedback on the simulation work. We would also like to thank Johanna Waalkens for her initial contribution in staining optimisations.

## Author contributions

Conceptualization: RK, TM

Methodology: RK, HK, EG, TR, MADRBFL, WM, TM

Investigation: RK, HK, EG, TM

Data analysis: RK, HK, MADRBFL

Visualization: RK, HK, EG, MADRBFL

Supervision: EG, FW, WM, TM

Writing—original draft: RK, TM

Writing—review & editing: RK, HK, EG, MADRBFL, TM

