## Supplementary Material and Figures for "Charting the nanotopography of inner hair cell synapses using MINFLUX nanoscopy"

### Supplementary Materials and Figures

#### Descriptions of alternative sample preparation approaches tested for MINFLUX imaging

##### HARD fixation

For HARD experiments (*33*), we used Ai32VC cre$^+$ mice (*105*) which express channel rhodopsin 2 (ChRh2) with a fluorescent EYFP tag under an IHC-specific VGLUT3 promoter. This approach allowed us to readily identify IHCs on coverslips. We adopted two approaches for heat-assisted immobilization of IHCs (Fig S1):

1. Cochleae were dissected in ice-cold PBS. The round window of the cochlea was injected with 35% Pluronic gel (Pluronic F127 NF; BASF, Parsippany, NJ) using a pre-cooled syringe as described before (*106*) to support partitioning during cochlear sectioning. The cochlea was glued using cyanoacrylate on the stage of a slicing chamber filled with PBS at room temperature which causes Pluronic Gel to solidify. We then used a Leica VT1000s Vibratome to obtain 200 µm thick sections (slicing parallel to the cochlear modiolus, minimum speed and maximum frequency setting). Slices were immediately transferred to a poly-L-lysine coated coverslip maintained at 50°C on a hot plate. We tested different time durations ranging from 30 seconds to 5 minutes. The excess tissue was removed, and the coverslips were allowed to cure on the hot plate for another 5 minutes.
2. Cochleae were dissected in ice-cold PBS and the apical end of the organ of Corti was further dissected out and the tectorial membrane was carefully removed using a pair of fine forceps. The acute preparation of the organ of Corti was placed directly on a poly-L-lysine coated coverslip heated to 50°C on a hot plate for different time durations ranging from 30 seconds to 5 minutes. The excess tissue was removed, and the coverslips were allowed to cure on the hot plate for another 5 minutes. We tried placing the organ of Corti top-up with the basilar membrane towards the coverslip as well as tried positioning it in an inverted manner with the cuticular plate towards the coverslip. Both approaches yielded immobilized layers of cells on the coverslip.

After either approach, coverslips were immunostained with anti-chicken GFP (Abcam, ab13970) and anti-mouse CtBP2 (BD Biosciences, 612044) as described before in the methods section. With the first approach, results were largely inconsistent for us and in very few instances (2 out of 7 trials) we could obtain identifiable individual IHCs with synaptic ribbons on the coverslip. On the other hand, the second approach appeared more promising, with rows of identifiable IHCs with synaptic ribbons (Fig. S1B, C) in all experimental trials (N = 3). While the signal:noise ratio appeared optimal, MINFLUX did not work for these samples largely on account of larger distance of the synapses from the coverslip (>3 µm). It appeared this was due to IHCs sticking to the coverslip with either the basilar membrane in between, or IHCs were anchored from their apical end with the cuticular plate at the coverslip and the synapses away from it. An illustration depicting the two possible scenarios has been provided in Fig S1D.

**Cochlear Cryosections**

Cryosections of cochleae from 2-week-old C57BL6/J mice were performed as described elsewhere (*79*). We collected the cryosections on poly-L-lysine coated coverslips (Fig S1E) and tested sections of varying thicknesses ranging from 10 µm to 16 µm and found comparable results (representative immunostained section shown in Fig S1F, G, H). While our initial attempts with MINFLUX imaging did not yield any results owing to high background fluorescence, treating the sections prior to blocking and immunostaining with 0.1% $NaBH_4$ solution (7 minutes) or 0.1M Tris (30 minutes) quenched tissue autofluorescence significantly. This allowed us to obtain some preliminary 2D (Fig S1I) and 3D (data not shown) MINFLUX images of synaptic ribbons (RIBEYE), with suboptimal imaging quality. MINFLUX imaging of any other synaptic protein labelling did not work.

#### Supplementary text for biophysical modelling

##### Estimation of the local calcium concentration around synaptic vesicle sensors

We placed 6 sensors around the synaptic vesicle (SV) structures. As the location of sensors is unknown, we first estimated the local [$Ca^{2+}$] around the SV (docked). To estimate local [$Ca^{2+}$] around sensors, we used "hits" as a proxy of the [$Ca^{2+}$] (*107*) where the number of hits represents the number of $Ca^{2+}$ collisions on the vesicle surfaced by MCell. At every time step of simulations, the number of hits and the location of the hit event are stored. As regions of the vesicle surface proximal to the nanocluster exhibited higher hit rates compared to distal regions, we divided the vesicle surface into 648 discrete surfaces to reflect the spatial dependency of the local calcium concentration. We calculated the density of hits per unit area instead of summating over the whole surface as follows:

$$unit\ area\ = r^2 \cdot \cos\theta \cdot\ \delta\theta\ \cdot \delta\varphi$$

where $\theta$ is a polar angle ($0\ \leq \theta \leq \pi$) and $\varphi$ is the azimuth angle ($-\pi \leq \varphi \leq \pi$). The polar angle was divided into 18 bins and the azimuth angle was divided into 36 bins. To convert the hit rates per unit area to the local calcium concentration, we ran simulations with one synaptic vesicle in a space filled with $Ca^{2+}$ ions with a concentration of 20, 50, 100 $\mu M$. From these simulations, we could calculate the hit rate per unit area for each concentration. Then, we selected four-unit areas near the sensor 1 – which triggered the majority of release events - and plotted the mean of four unit areas (Fig. S7B). We summed the hit rate per unit area from t = 1.5 ms to t = 2 ms where the hit rate reaches the plateau for the single nanocluster. Linear regression was used to estimate the local [$Ca^{2+}$] near sensor index 1, yielding an approximate value of 40 μM for the sensor index 1 in the single nanocluster case (hit rate per unit area ($\times 10^6$) = $5\times10^4$ [$Ca^{2+}$] + $15\times10^4$).

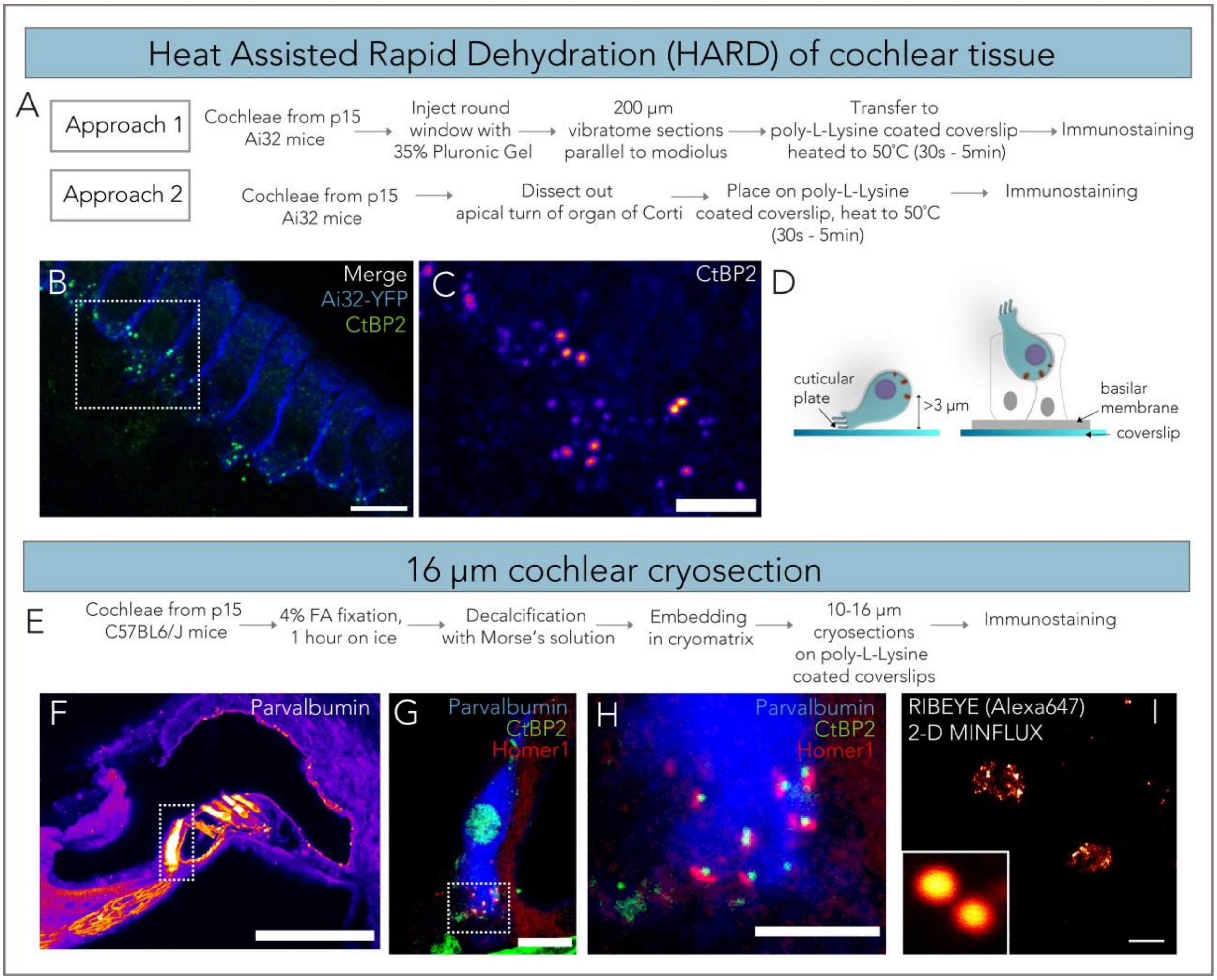


**Fig. S1. Summary of alternative attempts at sample preparation for MINFLUX of IHC ribbon synapses**

**(A)** Two alternative approaches towards Heat Assisted Rapid Dehydration (HARD) of cochlear tissue to fix IHCs on coverslip. Only approach 2 could be used for reproducible tissue immobilization on coverslips. **(B)** Confocal (single plane) image of HARD immobilized tissue from Ai32 mice showing IHC membrane (blue) and synaptic ribbons (CtBP2, green). Scale bar = 10 µm. **(C)** Confocal zoom-in showing synaptic ribbons in the region marked in (B). An intensity-coded look-up table has been used for depiction; scale bar = 5 µm. **(D)** IHCs appeared to be immobilized on the coverslips either via their apical end or via the basilar membrane, depending on the orientation of placement of the organ of Corti during HARD fixation. Either way, synapses were typically at a distance greater than 3 µm from the coverslip, making MINFLUX imaging difficult. **(E)** 10 – 16 µm cochlear cryosections for MINFLUX **(F)** Representative confocal overview of the organ of Corti from a 16 µm cochlear cryosection, scale bar = 100 µm. **(G, H)** Maximum-intensity projection of IHC marked in (F) shows ribbon synapses (CtBP2, green) and the postsynaptic density (Homer1, red). Scale bar = 10 and 5 µm for (G) and (H) respectively. **(I)** Exemplary 2D MINFLUX imaging of RIBEYE (Alexa 647) from a 16 µm cochlear cryosection. Inset shows corresponding confocal view of the two ribbons shown. Scale bar = 200 nm.

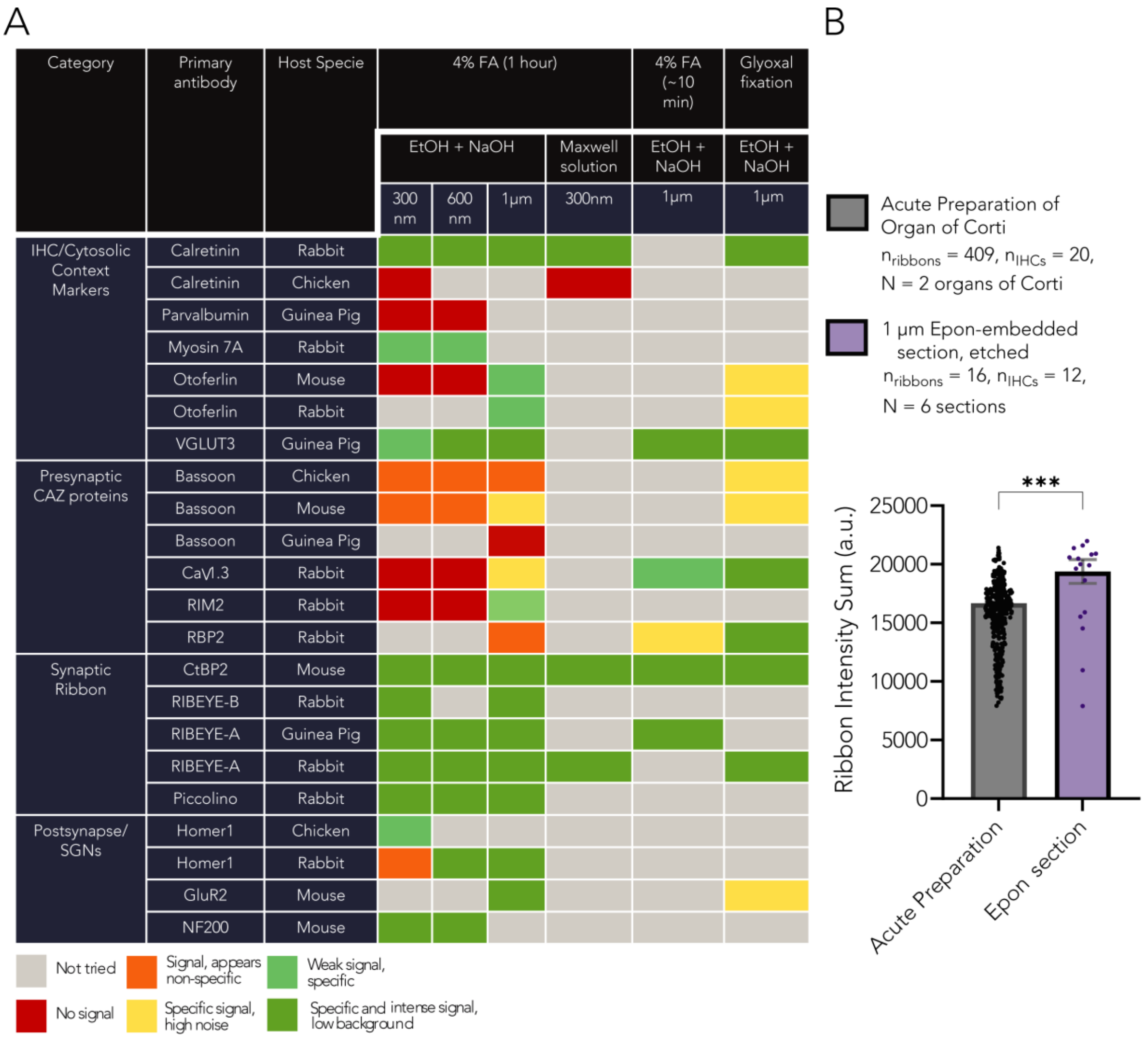


**Fig. S2. Optimization of immunostainings of semithin sections from Epon embedded cochleae after etching** **(A)** Summary of primary antibody stainings attempted on sections of varying thickness (300 nm, 600 nm and 1 μm) and the subjective quality of stainings have been denoted with a colour code. Several immunostainings that did not work for thinner sections showed drastic improvement on switching to 1 μm sections, possibly because of more intact synaptic structures. In addition to etching samples with NaOH and ethanol, we initially also performed etching of 300 nm thin sections with Maxwell solution (*108*) and found very comparable results. Immunolabeling for RBP2 and $Ca_V$1.3 showed poor results when cochleae were fixed with 4% formalin for 1 hour. The quality of stainings for these improved upon a shorter fixation duration for ~10 minutes and was the most optimal with glyoxal fixation. **(B)** Intensity of immunostained synaptic ribbons was compared between acute preparations of the organ of Corti and Epon-embedded and etched 1 μm sections. The samples were both from postnatal day 15 C57BL/6J mice, fixed for 1 hour with 4% formalin, decalcified with Morse's solution, were stained with the same antibody concentrations and imaged with the same laser settings in parallel. Epon embedding and etching the samples with NaOH and ethanol does not seem to reduce immunofluorescence signal intensity, suggesting that epitopes are preserved well through the process.

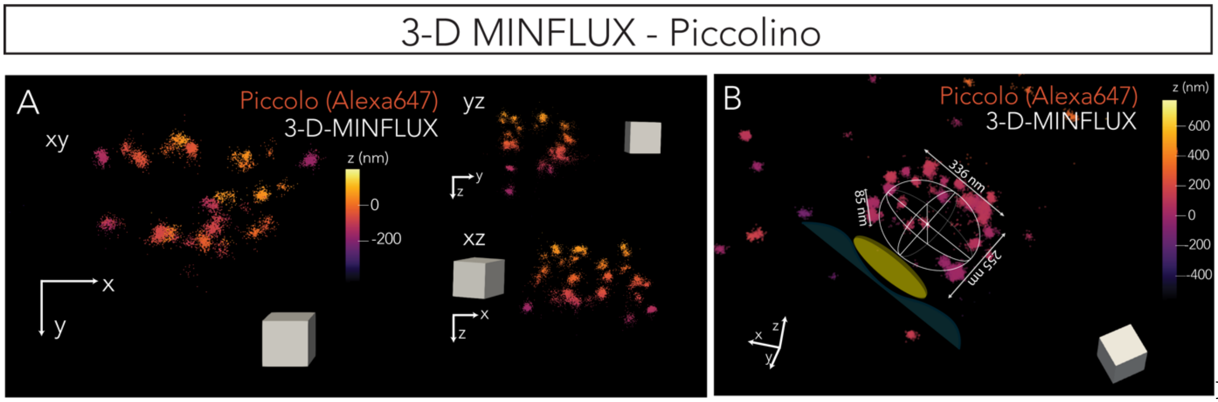


**Fig. S3. 3D MINFLUX images of Piccolino show an arch-shaped structure that envelops the synaptic ribbon**

**(A)** Representative 3D MINFLUX image of Piccolino (labelled with an antibody recognizing amino acid 2012–2351 of rat Piccolo), shows a characteristic arch shaped structure along the z-projection. Projections along the x- and y- axes have also been depicted. **(B)** Another exemplary 3D MINFLUX image of Piccolino from a different IHC. An illustrated ellipsoid showing the likely position of the synaptic ribbon inside the Piccolino arch has been overlaid. The image has been tilted as shown by the orientation axis in the left corner to better reveal the structure. Colour codes represent the distances along the z-axis; the cube shown for scale has an edge of length 100 nm.

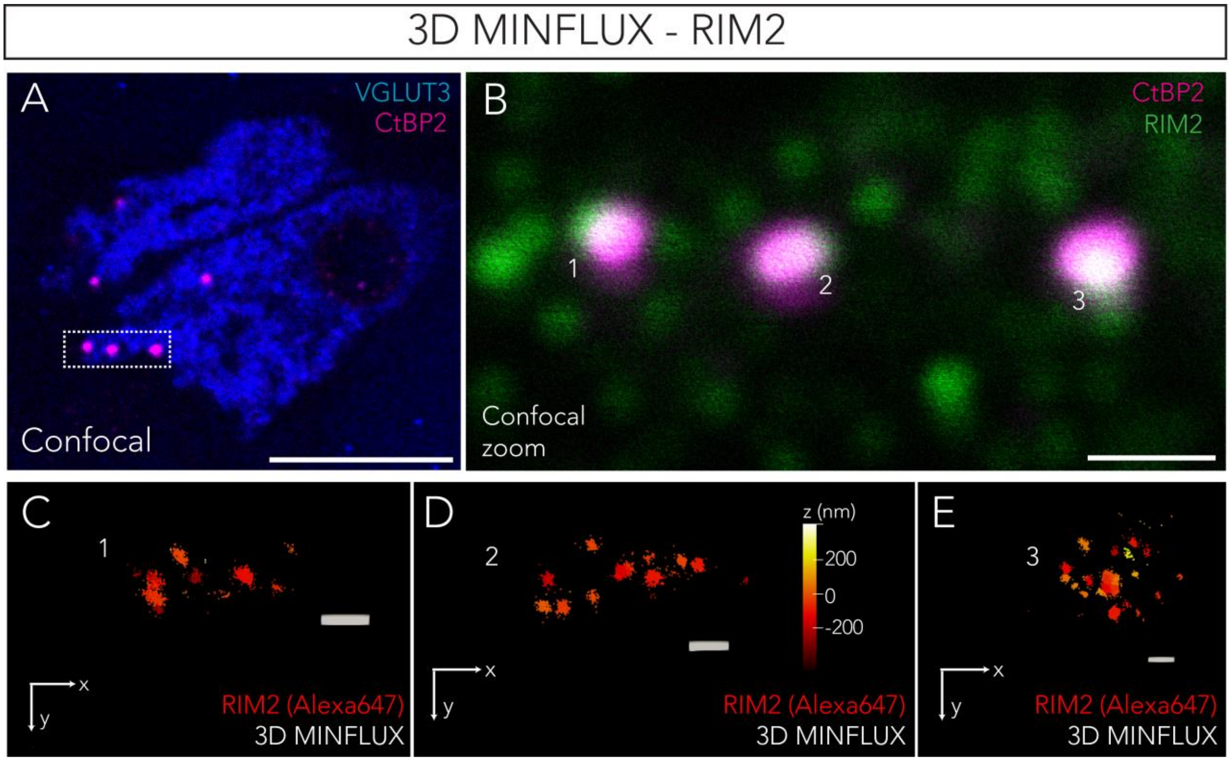


**Fig. S4. 3D MINFLUX images of AZ protein RIM2**

**(A)** Representative confocal overview (single plane) of IHC used for imaging (VGLUT3, blue) and the location of the ribbon synapses in the cell (CtBP2, magenta). Scale bar = 10 µm. **(B)** Zoom-in of the ribbon synapses. Here the corresponding RIM2 puncta have been imaged (green). Scale bar = 1 µm. **(C, D, E)** Z-projections of 3D MINFLUX images of RIM2 from the synapses marked in (B) shows the topographic distribution of the protein at the AZ. Colour code represents distances along the z-axis and scale bar has a length of 100 nm.

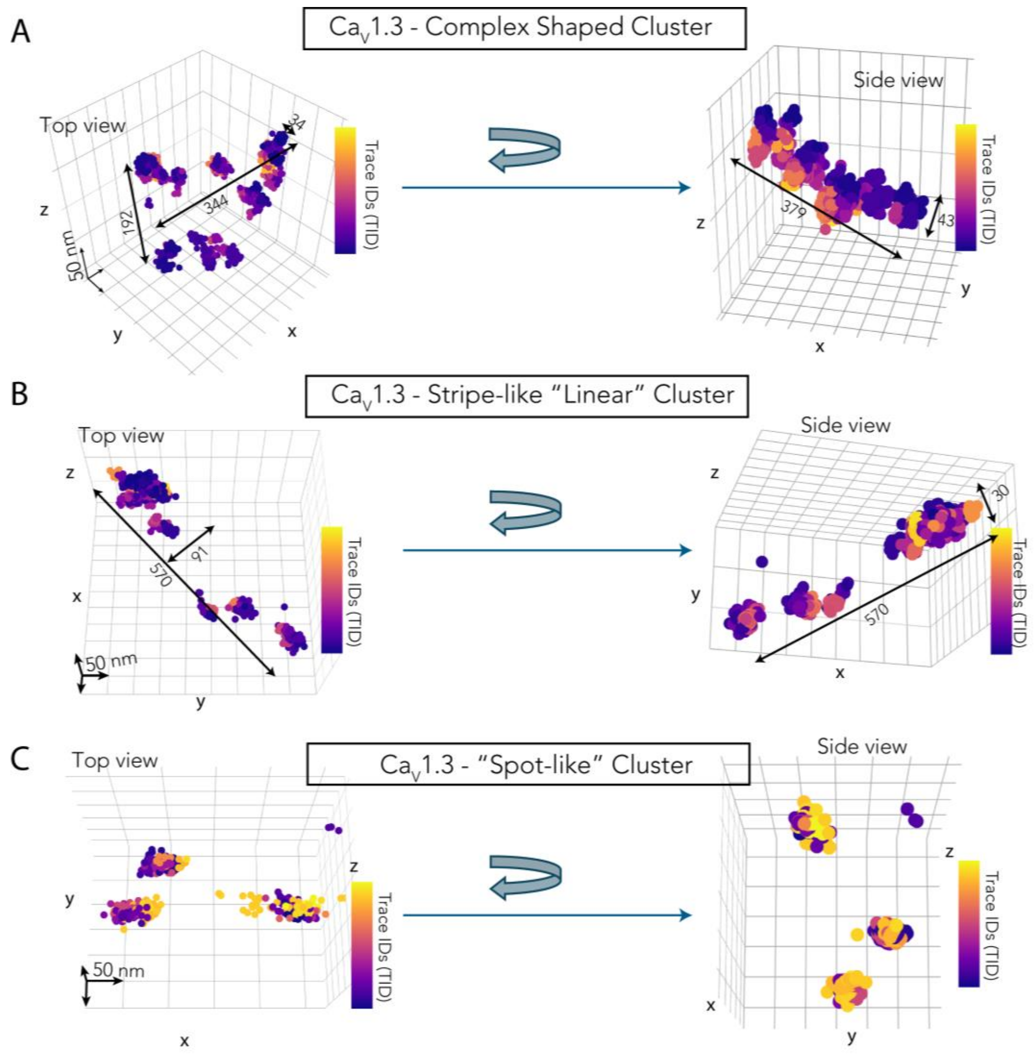


**Fig. S5. Heterogeneous morphology of Ca$_V$1.3 channel clusters in 3D MINFLUX images.** Further exemplary synapses (raw 3D MINFLUX data) illustrating the heterogeneity in morphology of Ca$_V$1.3 channel clusters observed in IHCs. **(A)** Distribution of Ca$_V$1.3 localisations shows a complex spatial arrangement when viewed from the top, however rotating the image displays a linear arrangement of the localisations along the side, likely presenting a view along the plasma membrane as also shown in Fig 2H. Such “complex” channel clusters account for 44.5% of synapses in our data. **(B)** A linear, stripe-like Ca$_V$1.3 cluster; the stripe-like appearance is evident irrespective of the orientation in which the synapse is viewed. Such synapses also make up 44.5% of the total synapses we imaged. **(C)** Small “spot-like” Ca$_V$1.3 cluster with no define morphology; these constitute the remaining 11% of synapses in our data. Representative measures of dimensions have been depicted with arrows; scale of grid lines is 50 nm along all dimensions and colour code represents Trace IDs of individual localisations.

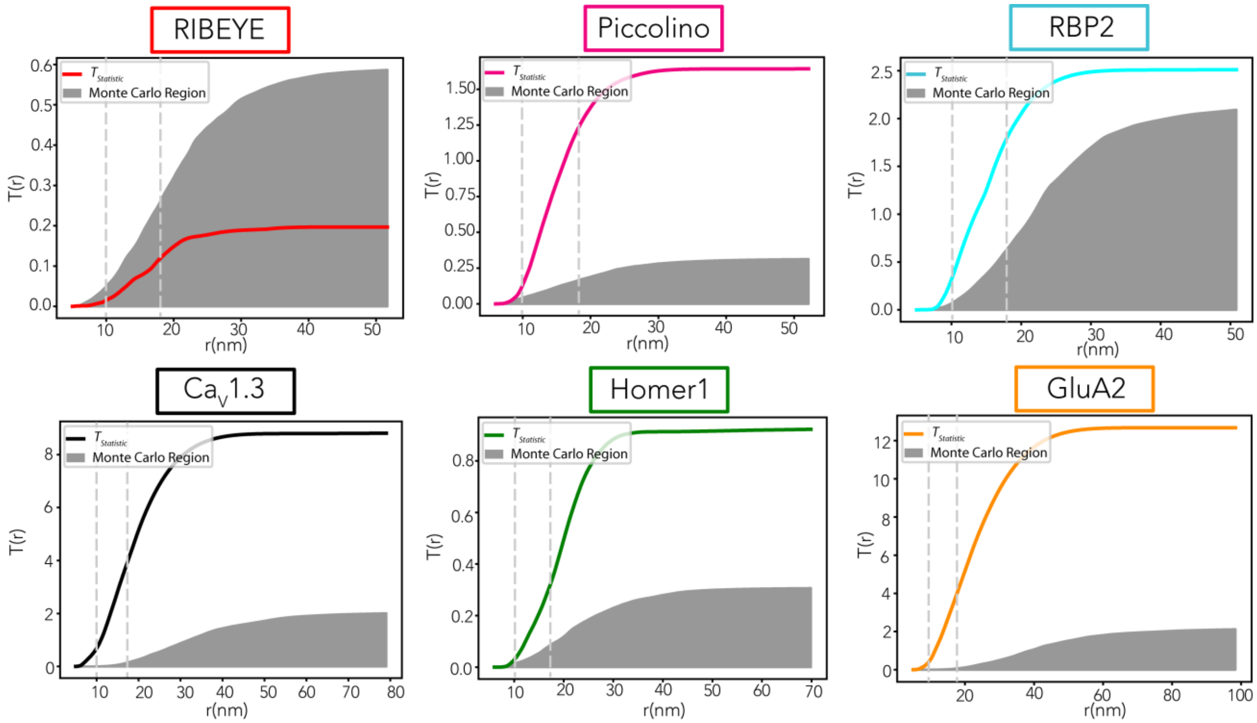


**Fig. S6. Envelop representation of DCLF Test**

Exemplary envelope representations for the Diggle-Cressie Loosmore-Ford (DCLF) test of spatial randomness (SR) confined to protein molecule's convex hull, applied to the data of Fig. 5, and considering the full range of distance values from experiment localization precision to tens of nanometres. Vertical lines represent the evaluation interval. DCLF test statistic T (solid-coloured lines) and Monte Carlo acceptance/non-rejection region (shaded) have been plotted as a function of the length r of the distance interval. Graphs based on 1000 simulations of SR.

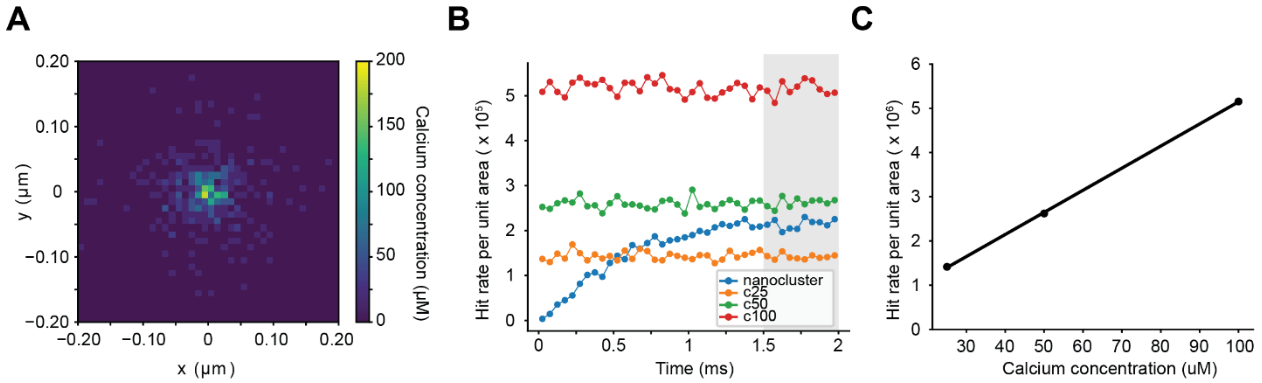


**Figure S7. Concentration estimation for a single nanocluster scenario.**

**(A)** Calcium concentration map of a single nanocluster. Each pixel represents a voxel with a size of 10 nm × 10 nm × 10 nm. The range of z is from 0 to 10 nm. Color bar indicates the range of calcium concentration (unit: µM). **(B)** The plot of hit rate per unit area in time for four selected surface regions located near sensor index 1. Each trace corresponds to the single nanocluster case (blue) and the conditions with fixed calcium concentrations of 25, 50, and 100 µM (orange, green, red, respectively). The shaded grey region (1.5 – 2.0 ms) indicates the plateau phase in the single nanocluster case (blue). **(C)** The fitted curve showing the relationship between the calcium concentration and the summation of the hit rate per unit area within the time window of 1.5 – 2.0 ms.

#### Table S1. List of primary antibodies

| Antibody | Host specie | Epitope | Dilution | Company | Identifier |
|---|---|---|---|---|---|
| Anti-Calretinin | Rabbit Polyclonal | - | 1:200 | Swant | CR7697 |
| Anti-Calretinin | Chicken Polyclonal | Full length recombinant mouse Calretinin (UniProt ID: Q08331) | 1:200 | Synaptic Systems | 214106 |
| Anti-Parvalbumin | Guinea Pig Polyclonal | - | 1:200 | Synaptic Systems | 195004 |
| Anti-MyoVIIA | Rabbit Polyclonal | AA 880 – 1077 of tail region of human myosin-VIIa | 1:200 | Proteus | 25-6790-C050 |
| Anti-Otoferlin | Mouse Monoclonal | AA 1 – 400 of recombinant human Otoferlin | 1:200 | Abcam | ab53233 |
| Anti-Otoferlin | Rabbit Polyclonal | AA 181 – 196 of mouse Otoferlin (UniProt ID: Q9ESF1-1) | 1:200 | Synaptic Systems | 178003 |
| Anti-VGLUT3 | Guinea Pig Polyclonal | AA 543 – 601 of mouse VGLUT3 (UniProt ID Q8BFU8) | 1:200 | Synaptic Systems | 135204 |
| Anti-$Ca_V1.3$ | Rabbit Polyclonal | AA 859 – 875 of rat $Ca_V1.3$ (Accession P27732), intracellular loop between domains II and III | 1:100 | Alomone | ACC-005 |
| Anti-RIM2 | Rabbit Polyclonal | AA 461 – 987 of rat RIM2 (PDZ domain) | 1:200 | Synaptic Systems | 140103 |
| Anti-RBP2 | Rabbit Polyclonal | AA 596 – 868 of mouse RBP2 (UniProt ID Q80U40) | 1:200 | Synaptic Systems | 316103 |
| Anti-CtBP2 | Mouse Monoclonal IgG1 | Mouse CtBP2 AA 361 – 445 | 1:200 | BD Biosciences | 612044 |
| Anti-RIBEYE-B | Rabbit Polyclonal | AA 974 – 988 of rat RIBEYE (UniProt ID Q9EQH5-2) | 1:200 | Synaptic Systems | 192003 |
| Anti-RIBEYE-A | Rabbit Polyclonal | AA 95 – 207 of rat RIBEYE (UniProt ID Q9EQH5-2) | 1:200 | Synaptic Systems | 192103 |
| Anti-RIBEYE-A | Guinea Pig Polyclonal | AA 95 – 207 of rat RIBEYE (UniProt ID Q9EQH5-2) | 1:200 | Synaptic Systems | 192104 |
| Anti-Piccolo | Rabbit Polyclonal | AA 2011 – 2350 of rat Piccolo (UniProt ID Q9JKS6) | 1:200 | Synaptic Systems | 142113 |
| Anti-Bassoon | Chicken Polyclonal | Recombinant protein corresponding to central region rat Bassoon (UniProt Id: O88778) | 1:200 | Synaptic Systems | 141016 |
| Anti-Bassoon | Guinea Pig Polyclonal | Recombinant protein corresponding to central region rat Bassoon (UniProt Id: O88778) | 1:200 | Synaptic Systems | 141004 |
| Anti-Bassoon | Mouse Monoclonal IgG2a | AA 756 – 1001 of mouse Bassoon (clone SAP7F407) | 1:200 | Abcam | Ab82958 |
| Anti-Homer1 | Rabbit Polyclonal | N-terminal of human Homer1 (UniProt ID Q86YM7) | 1:200 | Synaptic Systems | 160002 |
| Anti-Homer1 | Chicken Polyclonal | Recombinant protein corresponding to N-terminal | 1:200 | Synaptic Systems | 160006 |

| | | | | | |
|---|---|---|---|---|---|
| | | of human Homer1 (UniProt ID: Q86YM7) | | | |
| Anti-GluA2 | Mouse Monoclonal IgG2a | GluA2 extracellular (putative N-terminal portion, AA 175 – 430), UniProt ID P19491 | 1:200 | Merck Millipore | MAB397 |
| Anti-NF200 | Mouse Monoclonal IgG1 | Neurofilament Heavy Chain (200 kDa) clone NE14 | 1:200 | Sigma Aldrich | N5389 |

**Table S2. List of secondary antibodies**

| Antibody | Host specie | Dilution | Company | Identifier |
|---|---|---|---|---|
| Alexa 488 conjugated anti-rabbit | Goat IgG (H+L) | 1:200 | Invitrogen | A11008 |
| Alexa 488 conjugated anti-chicken | Goat IgG (H+L) | 1:200 | Invitrogen | A11039 |
| Alexa 488 conjugated anti-guinea pig | Goat IgG (H+L) | 1:200 | Invitrogen | A11073 |
| Alexa 488 conjugated anti-mouse | Goat IgG (H+L) | 1:200 | Invitrogen | A11001 |
| Alexa 546 conjugated anti-mouse | Goat IgG (H+L) | 1:200 | Invitrogen | A11003 |
| Alexa 647 conjugated anti-rabbit | Goat IgG (H+L) | 1:200 | Invitrogen | A21244 |
| Alexa 647 conjugated anti-mouse | Goat IgG (H+L) | 1:200 | Invitrogen | A21236 |
| Alexa 647 conjugated anti-mouse | Goat IgG2a | 1:200 | Invitrogen | A21241 |
| Alexa 647 conjugated anti-chicken | Goat IgG (H+L) | 1:200 | Invitrogen | A21449 |
| CF660C conjugated anti-mouse | Goat IgG (H+L) | 1:200 | Biotium | 20052-1 |
| CF660C conjugated anti-guinea pig | Goat IgG (H+L) | 1:200 | Biotium | 20497-1 |

**Table S3. 2D MINFLUX sequence parameters**

| Iteration | L (nm) | Photon threshold | CFR threshold | Dwell time (ms) | Pattern Repeats | Background offset (kHz) | Power factor |
|---|---|---|---|---|---|---|---|
| 0 | 288 | 160 | -1 | 1 | 1 | 15 | 1 |
| 1 | 288 | 150 | -1 | 1 | 5 | 10 | 1 |
| 2 | 151 | 100 | 0.8 | 1 | 5 | 10 | 2 |
| 3 | 76 | 100 | 0.8 | 1 | 5 | 10 | 4 |
| 4 | 40 | 150 | * | 1 | 5 | 10 | 6 |

*= checked, no threshold applied; CFR = Centre Frequency Ratio

**Table S4. 3D MINFLUX sequence parameters**

| Iteration | L (nm) | Photon threshold | CFR threshold | Dwell time (ms) | Pattern Repeats | Background offset (kHz) | Power factor |
|---|---|---|---|---|---|---|---|
| 0 | 288 | 160 | -1 | 1 | 1 | 15 | 1 |
| 1 | 288 | 400 | -1 | 1 | 1 | 15 | 1 |
| 2 | 288 | 100 | -1 | 1 | 5 | 10 | 1 |
| 3 | 288 | 50 | -1 | 1 | 5 | 10 | 1 |
| 4 | 151 | 67 | 0.9 | 1 | 5 | 10 | 2 |
| 5 | 151 | 33 | -1 | 1 | 5 | 10 | 2 |
| 6 | 76 | 67 | 0.8 | 1 | 5 | 10 | 4 |
| 7 | 76 | 33 | -1 | 1 | 5 | 10 | 4 |
| 8 | 40 | 100 | -1 | 1 | 5 | 10 | 6 |
| 9 | 40 | 50 | -1 | 1 | 5 | 10 | 6 |

CFR = Centre Frequency Ratio

#### Table S5. Summary of different morphologies of $Ca_V$1.3 clusters and quantified parameters

$Ca_V$1.3 clusters were classified into “lines”, “complexes” or “spot-like” as also depicted in Fig. S5. For tapered synapses (see Fig S5A), where one end appeared broader than the other end, we measured the width along the two ends and reported the average. For two “complex” and all “spot-like” synapses, measuring the three dimensions was not possible due to the irregular shape of the synapse (denoted as not applicable, “NA” in the table below). For four synapses (all lines), DCLF test indicated non-clustered spatial arrangement of molecular counts. For these synapses, nanocluster parameters were not quantified (denoted with a ‘-’).

| N | Morphology | Length (nm) | Width (nm) | Height (nm) | Mol. Counts | Alpha Shape Volume (µm$^3$) | No. of nano clusters | Average no. of channels/ nanocluster | Average Inter-cluster NND (nm) | Average Intra-cluster NND (nm) |
|---|---|---|---|---|---|---|---|---|---|---|
| 1 | line | 673.5 | 156.6 | 61.3 | 24.0 | 4.0E-04 | - | - | - | - |
| 2 | line | 359.4 | 152.7 | 69.7 | 42.0 | 1.3E-03 | 6.0 | 6.7 | 100.3 | 12.7 |
| 3 | line | 224.1 | 84.3 | 49.9 | 41.0 | 2.7E-04 | - | - | - | - |
| 4 | line | 414.2 | 97.3 | 55.3 | 28.0 | 9.0E-04 | - | - | - | - |
| 5 | line | 565.5 | 91.4 | 29.9 | 57.0 | 6.3E-04 | 7.0 | 7.1 | 64.0 | 8.6 |
| 6 | line | 455.6 | 84.1 | 39.7 | 51.0 | 2.0E-03 | 5.0 | 9.8 | 103.1 | 9.0 |
| 7 | line | 441.5 | 114.6 | 53.8 | 27.0 | 8.5E-04 | 2.0 | 10.5 | 131.8 | 13.8 |
| 8 | line | 241.1 | 38.3 | 20.2 | 12.0 | 6.0E-05 | - | - | - | - |
| **Average (lines only)** | | **421.9** | **102.4** | **47.5** | **35.3** | **8.0E-04** | **5.0** | **8.5** | **99.8** | **11.0** |
| 9 | complex | 479.6 | 190.6 | 52.3 | 43.0 | 2.1E-03 | 4.0 | 9.8 | 211.9 | 10.4 |
| 10 | complex | 267.5 | 174.6 | 57.6 | 66.0 | 3.1E-03 | 5.0 | 12.2 | 103.9 | 9.3 |
| 11 | complex | 388.6 | 146.9 | 43.1 | 70.0 | 1.6E-03 | 7.0 | 8.3 | 84.3 | 10.1 |
| 12 | complex | 484.9 | 156.6 | 48.2 | 40.0 | 2.5E-03 | 5.0 | 6.4 | 84.8 | 10.6 |
| 13 | complex | NA | NA | NA | 17.0 | 1.4E-03 | 3.0 | 4.0 | 195.7 | 14.0 |
| 14 | complex | 466.0 | 266.0 | 40.1 | 45.0 | 2.1E-03 | 8.0 | 4.8 | 114.5 | 11.9 |
| 15 | complex | 253.9 | 128.2 | 19.4 | 9.0 | 9.9E-05 | 2.0 | 4.0 | 215.6 | 8.4 |
| 16 | complex | NA | NA | NA | 23.0 | 1.5E-03 | 5.0 | 4.4 | 117.8 | 12.2 |
| **Average (complexes only)** | | **390.1** | **177.1** | **43.4** | **39.1** | **1.8E-03** | **4.9** | **6.7** | **141.1** | **10.9** |
| 17 | spot-like | NA | NA | NA | 17.0 | 6.5E-03 | 3.0 | 4.7 | 232.9 | 28.7 |
| 18 | spot-like | NA | NA | NA | 23.0 | 5.1E-04 | 3.0 | 7.0 | 140.3 | 9.3 |
| **Average (spot-like only)** | | **NA** | **NA** | **NA** | **20.0** | **3.5E-03** | **3.0** | **5.8** | **186.6** | **19.0** |
| **Average (all)** | | **408.2** | **134.4** | **45.7** | **35.3** | **1.5E-03** | **4.6** | **7.1** | **135.8** | **12.1** |

#### Table S6. Parameters for biophysical modelling

| **Parameter** | **Value** | | | | **Reference** |
|---|---|---|---|---|---|
| | Concentration | Diffusion Constants $[\mu m^2 ms^{-1}]$ | $K_D$ $[\mu\mathrm{M}]$ | $k_{on}$ $[\mu M^{-1} ms^{-1}]$ | |
| Fixed Buffers | 610 µM | | 4.859 | 1.375 | (*109*) |
| EGTA | 800 µM | 0.14 | 0.071 | 0.0105 | (*110*, *111*) |
| BAPTA | 400 µM | 0.14 | 0.17 | 0.45 | (*110*, *112*) |
| ATP | 68 µM (Calculated in 3mM $Mg^{2+}$ 2mM ATP) | 0.14 | 2200 | 0.013 | (*113*) |
| Resting $Ca^{2+}$ concentration | 50 nM | 0.223 | | | (*91*, *114*) |
